# Benchmarking strain-level profiling of *Escherichia coli* in short-read gut metagenomes

**DOI:** 10.64898/2026.05.19.726160

**Authors:** Matthew Galbraith, David Williams, Liam P. Shaw, Samuel Lipworth, Nicole Stoesser

## Abstract

2.

Metagenomes facilitate *Escherichia coli* strain-level profiling in human gut microbiomes, informing our understanding of colonisation diversity and the genetic features distinguishing infection from carriage. Among numerous reference-based tools for short-read metagenomic strain-level profiling, the best approach remains unclear. Here, we benchmarked six publicly available tools—PanTax, PathoScope 2.0, StrainGE, Strainify, StrainR2 and StrainScan—for their ability to detect co-existing strains of *E. coli* and estimate their relative abundance across real and simulated metagenomes of increasing complexity with varying reference database composition. We show that the tools varied substantially in both the number and relative abundance of strains predicted, indicating that despite being designed for the same broad purpose, they produce markedly different representations of strain composition from the same metagenomic samples. StrainR2 achieved the highest F1 score across our experiments (0.995) and successfully distinguished all strains within a complex, 10-strain spike-in mixture containing strains sharing up to 99.9% average nucleotide identity and included in a 304-strain *E. coli* reference database. However, when ground-truth strains were removed from the reference database to mimic the profiling of samples of unknown strain content PathoScope 2.0 and StrainGE performed strongest: both estimated phylogroup abundances accurately and frequently correctly assigned strain identities to the closest available genomic neighbours to the ground truth. Computational requirements also varied substantially (median runtime: 0.11– 68.45 h; peak memory: 2.08–351.53 GiB). Collectively, these results demonstrate that tool selection should reflect the intended application, and we provide practical guidance on the relative strengths and use cases of each approach. These findings will facilitate characterisation of *E. coli* strain-level diversity within short-read gut metagenomes with greater accuracy than previously possible.

**Impact statement:** Strain-level diversity within the human gut microbiome can be important for human health, with species such as *Escherichia coli* existing as both commensal and pathogenic strains. Most gut microbiome datasets are from short-read *i.e.*, Illumina, sequencing, and numerous bioinformatic tools have been developed to profile strain-level variation from these data. However, available tools have been benchmarked in various ways and are subject to author bias. This is, to our knowledge, the first independent benchmarking of six publicly available tools for *E. coli* strain profiling in short-read metagenomes. Using both real and simulated datasets of increasing complexity, we demonstrate tool performance varies in terms of strain detection and relative abundance estimation, highlighting that tool choice should be guided by the specific research question, as no single method performs optimally across all scenarios. This work provides an unbiased framework for tool selection, supporting more accurate and reproducible *E. coli* strain-level analyses in gut microbiome research from short-metagenomic data.

**Data summary:** The authors confirm all supporting data, code and protocols have been provided. Supplementary methods, tables and figures are available in the online Supplementary Material. Code for simulating metagenomes and generating *E. coli* spike-ins using InSilicoSeq, running tools on the simulated gut metagenomes and real-world spike-in datasets, inclusive of computational benchmarking, and R visualization and statistical analysis are available within a dedicated GitHub repository (https://github.com/mattgal11/benchmarking_short_read_strain_profilers). The following supplementary data files required for reproducible analyses are available on FigShare (https://doi.org/10.6084/m9.figshare.32125474):

- ZymoBIOMICS® D6331 gut microbiome standard dataset predicted relative abundances (Zymobiomics_D6331_raw_predicted_abundance.csv)
- SRR13355226 mock community paired-end reads with human reads depleted (SRR13355226_depleted_R1.fastq.gz & SRR13355226_depleted_R2.fastq.gz)
- SRR13355226 mock community dataset predicted abundance data, with and without human read removal (SRR13355226_raw_predicted_abundance_with_and_without_human_read_r emoval.csv)
- Normalised per-contig relative abundances for 98 species used to construct the baseline gut microbiome profile for InSilicoSeq metagenome simulation (Normalised_relative_abundance_for_InSilicoSeq_simulated_metagenomes_ gut_microbiome_profile.csv)
- Simulated metagenomes dataset call types and detection metric across detection thresholds (Simulated_metagenomes_raw_call_type_assingments_and_detection_thres holds.csv)
- Simulated metagenomes dataset (all references) predicted relative abundances (Simulated_metagenomes_all_references_raw_predicted_abundances.csv)
- Simulated metagenomes dataset (all references) mapped reads for PathoScope and Strainify (all_refs_pathoscope_reads_mapped.csv & all_refs_strainify_reads_mapped.csv)
- Simulated metagenomes dataset (reduced reference database) predicted relative abundances (Simulated_metagenomes_K12_and_Sakai_removed_from_reference_datab ase_raw_predicted_abundance.csv)
- *E. coli* reference genomes used in real-world spike ins dataset (304_ecoli_strains_reference_genomes.csv)
- NCBI datasets assembly stats data frame for 304 *E. coli* reference genomes (abram_et_al_assembly_stats.xlsx)
- FastANI all-vs-all whole-genome ANI for 304 *E. coli* reference database (fastANI_dRep_genomes_all_vs_all_real_world_spike_ins.tsv)
- Real world spike-ins dataset call types and detection metric across detection thresholds (real_world_spike_ins_detection_metrics_all_thresholds.csv)
- Real world spike-ins dataset predicted relative abundances (real_world_spike_ins_benchmarking_results.csv)
- Real world spike-ins dataset (reduced reference database) predicted relative abundances (real_world_spike_ins_benchmarking_no_ground_truth_results.csv)
- Real world spike-ins dataset computational benchmarking results (real_world_spike_ins_comp_benchmark_DB_step.csv & real_world_spike_ins_comp_benchmark_profiling.csv & strainR2_memory_comparisons.csv)
- Real world spike-ins dataset paired metagenome file sizes (real_world_spike_ins_metagenome_file_sizes.csv)

## 5. Introduction

The human gut microbiome is highly diverse and comprises >270 species[1], with substantial inter-individual variation that has been identified as a critical determinant of human health[2]. However, gut microbiome variation extends beyond species to the strain-level, with strains of the same species exhibiting genotypic and phenotypic heterogeneity[3, 4]. One example is *Escherichia coli*, with recent studies reporting an average of ∼2.5 unique strains in the gut per individual and dynamic strain turnover over time[5–8]. While commensal *E. coli* strains contribute to gut homeostasis[9, 10], extraintestinal pathogenic *E. coli* lineages contribute to both colonisation and disease and are the major cause of Gram-negative bloodstream infections worldwide[11–14]. Accurately characterising *E. coli* colonisation diversity in the gut microbiome would facilitate an improved understanding of the distribution and differences between commensal and disease-associated strains, informing targeted interventions such as vaccines and therapeutics.

Metagenomic sequencing has become the dominant method for gut microbiome analyses[15]. However, analysing *E. coli* strain-level diversity within available gut metagenomes remains challenging due to low relative abundances (typically ∼0.1– 1.2% in healthy individuals)[7, 16] and the historic predominance of short-read Illumina sequencing, which is limited in resolving repetitive genomic regions and consequently constrains the resolution of commonly used taxonomic classifiers[17]. For example, Kraken2 relies on shared *k*-mer matches and lowest common ancestor assignment, such that reads mapping to multiple closely related reference genomes are often classified at higher taxonomic levels[18, 19]. Similarly, *E. coli*-specific metagenomic typing approaches such as metaMLST infer sequence types using multi-locus sequence typing (MLST), but only to a single best-supported allele[20]. This can limit the ability to capture within-sample allelic diversity and restricts inference to dominant strains, as was seen within a recent analysis of 5,128 healthy adult gut metagenomes[21].

To address these limitations, numerous strain-level profiling tools for short-read metagenomes have been developed, employing different methodologies. However, the available tools have been benchmarked in various ways making direct comparisons difficult. Here, we survey existing tools, selecting six for further investigation using pre-defined criteria. We then benchmark their ability to profile *E. coli* strains across real and simulated metagenomes of increasing complexity, allowing us to provide recommendations for tool selection depending on the biological question.

## 6. Methods

### Selection of strain-level profiling tools

Candidate strain-level profiling tools were identified through literature searches in PubMed (https://pubmed.ncbi.nlm.nih.gov) and by examining references and citation networks of relevant publications. Tools were included in the benchmark if they (i) supported multi-strain-level resolution from short-read metagenomes, (ii) reported strain-level relative abundances, (iii) accepted a user-customisable reference database, and (iv) were actively maintained in an open-source repository *e.g.*, GitHub (Table 1). Tools which did not satisfy these requirements are listed in Table S1, alongside exclusion rationale. The six tools used were StrainScan[22] v1.0.14, Strainify[23] v1.3.1, StrainR2[24] v2.3.0, PathoScope 2.0[25] v2.0.7, PanTax[26] v2.1.0 and StrainGE[27] v1.3.9. For clarity when reporting numerical results, PathoScope 2.0 is hereafter referred to as PathoScope.

**Table 1.**
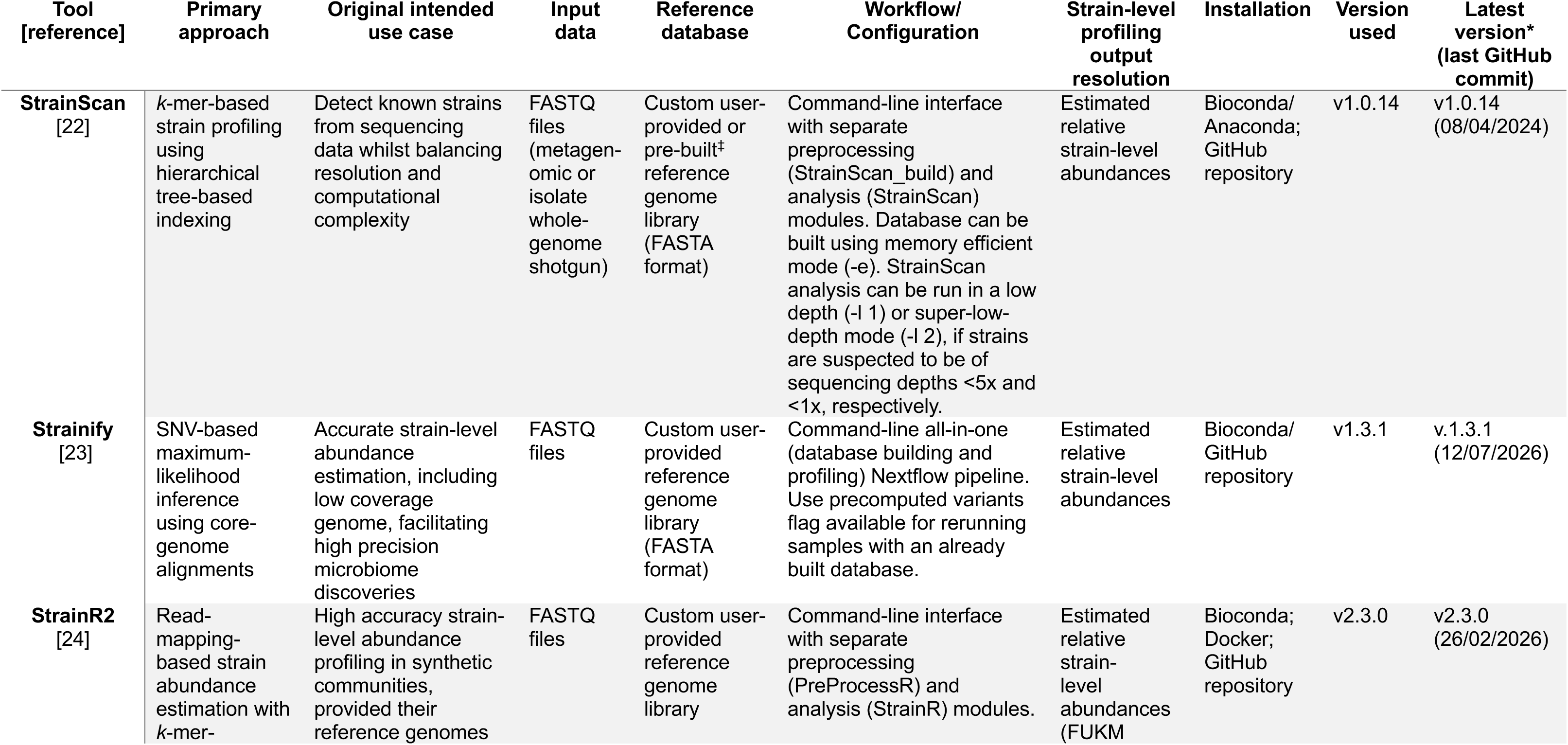
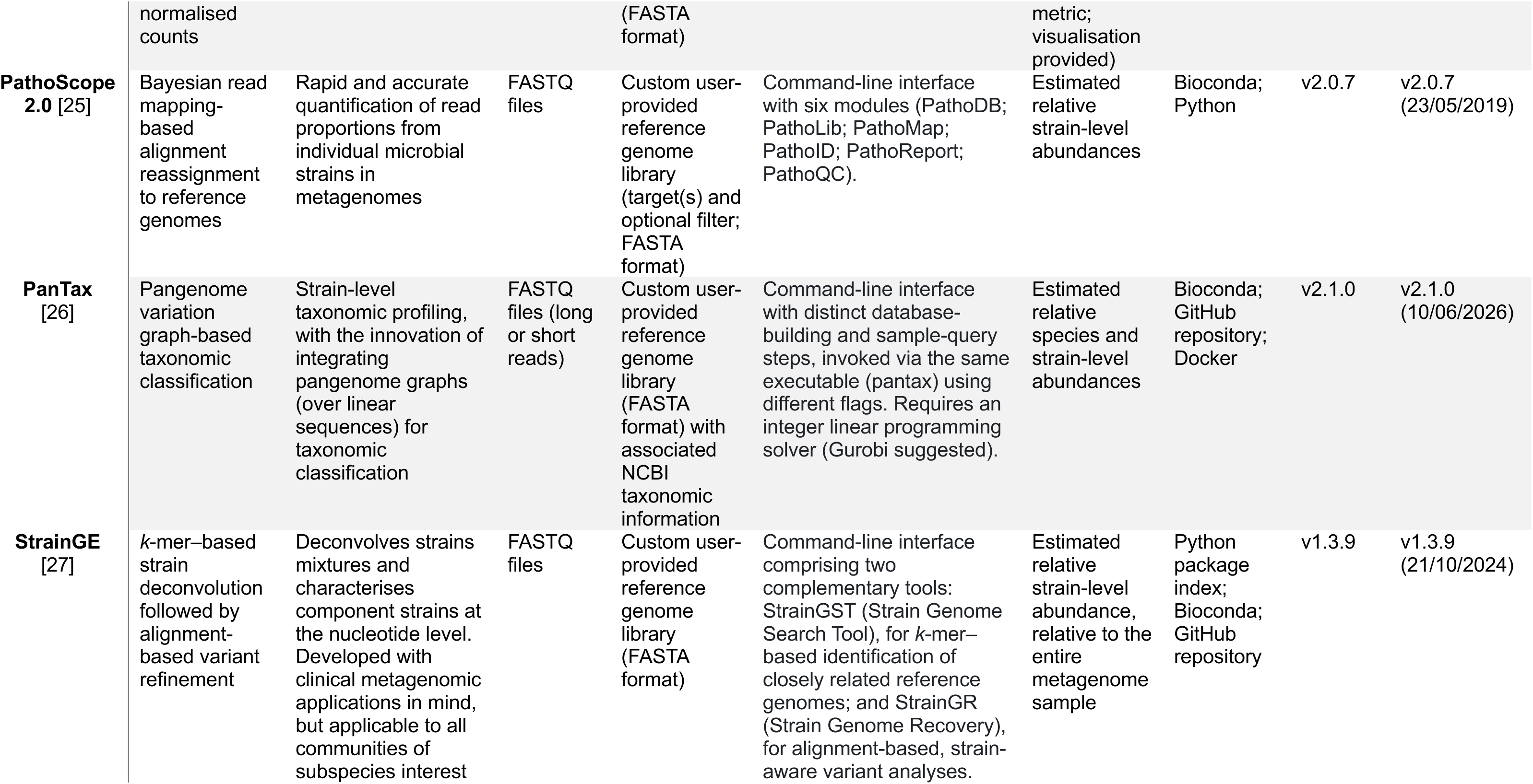
Reference-based, short-read metagenome strain-level profiling tools benchmarked in this study. *Most recent version as of 14/09/2026. ^‡^Pre-built databases for *Akkermansia muciniphila, Cutibacterium acnes, Prevotella copri, Escherichia coli, Mycobacterium tuberculosis, Staphylococcus epidermidis, Staphylococcus aureus* and *Lactobacillus crispatus*

### Computational environments

Tools and their dependencies were installed in dedicated conda environments using bioconda and conda-forge. StrainR2 required BBMap[28] v39.26 instead of v39.52 due to a bug affecting compatibility. Tools were executed as recommended, unless otherwise specified. Example commands used to run each tool are available (Supplementary Methods: S1).

### Benchmarking datasets

Across benchmarking datasets, default outputs for predicted relative abundance were taken per tool, unless otherwise stated. StrainGE was an exception, where estimated strain-level abundances from the StrainGST module (*rapct*; reported relative to the entire metagenome) were normalised (*rapct*/total_*rapct*) to enable comparisons.

Benchmarking datasets were designed to differentially test tools by evaluating: (i) different *E. coli* strains present at equal abundance in a low-complexity polymicrobial background (ZymoBIOMICS® D6331 gut microbiome standard), (ii) the impact of human read removal and differential abundance on *E. coli* strain profiling (SRR13355226 mock community), (iii) profiling of two differentially abundant *E. coli* strains present at differential sequencing depths in a polymicrobial background (simulated gut metagenomes), and (iv) the impact of increasing genomic similarity (%average nucleotide identity (ANI)) and differential relative abundance in profiling multiple *E. coli* strains spiked into publicly available, human gut metagenomes depleted of *E. coli* reads (real-world spike-ins). A brief description of each benchmarking dataset and the inputs provided to each tool is given below, with additional details on dataset construction and processing provided in the online Supplementary Material (see Supplementary Methods: S2).

#### (i) ZymoBIOMICS® D6331 gut microbiome standard

The ZymoBIOMICS® D6331 gut microbiome standard (Zymo Research, USA) contains 21 microbial strains, including 5 *E. coli* strains (JM109, B-3008, B-2207, B-766, B-1009) present at equal relative abundance (2.8%; Table S2). The downloaded short-read dataset (ShortPE_mock; SRR25211204) was provided as input for all tools, alongside a reference genome database of the five D6331 standard *E. coli* strains (Table S2). For PathoScope, initial relative abundance estimates were used instead of Bayesian-redistributed final guess estimates, as the latter assume unequal genome abundances, inconsistent with the dataset.

#### (ii) SRR13355226 mock community

SRR13355226 is a paired-end Illumina sequencing run (total sequencing output: 6.48 Gb) derived from a mock community consisting of 99% human DNA and 1% *E. coli*DNA (w/w), originally described in the StrainGE paper[27]. The 1% *E. coli* DNA comprises four strains (H10407, UTI89, O157:H7 str. Sakai, O139:H28 str. E24377A) at differing relative abundances, with the estimated sequencing depth for each strain also being calculated (Table S3). SRR13355226 and SRR13355226 depleted of human reads were used as FASTQ input to all tools, alongside a reference genome database comprising the four SRR13355226 mock community *E. coli* strains (Table S3). For this and all subsequent analyses, PathoScope final guess relative abundance estimates were used, whilst StrainScan was also run in super-low-depth mode (-l 2) with probabilistic strain detection enabled (-b 1).

#### (iii) Simulated gut metagenomes

We simulated gut metagenomes reflecting a realistic baseline healthy adult gut microbiome, informed by GutFeelingKB; a publicly available reference microbiome and species-level abundance profile[29]. In brief, 98 species were included in the baseline profile. *E. coli* was present at a relative abundance of 0.0196. This was split between the NCBI RefSeq strains (K-12 substr. MG1655 (GCF_000005845.2) and O157:H7 str. Sakai (GCF_000008865.2)) across a range of relative abundance ratios, inclusive of a negative control (Supplementary Methods S2; Table S4). Gut metagenomes were simulated across various sequencing depths using InSilicoSeq[30] v2.0.1 (Table S4).

To increase reference database complexity, a 24-strain *E. coli* reference database was constructed by selecting three representative genomes per major phylogroup (Table S5), with pairwise ANI comparisons calculated (Table S6). The 24 *E. coli* strain reference database and simulated metagenomes were used as input for all tools. To assess the impact of not including the strains present in the metagenome in the reference database, analyses were repeated using a reference database excluding *E. coli* K-12 substr. MG1655 and O157:H7 str. Sakai, leaving 22 strains. Outputs were averaged across triplicates for both experiments.

#### (iv) Real-world spike-ins

To evaluate tool performance under realistic community complexity with increased *E. coli* diversity, we performed a 10-strain spike-in experiment using publicly available metagenomes. In brief, 42 healthy adult short-read gut metagenomes spanning two studies (SRP373424/ERP005534; Table S7) were used[21]. These were depleted of *E. coli* reads using Kraken2[18] v2.17.1 and the extract_reads.py script from KrakenTools[33] v1.2.1 (https://github.com/jenniferlu717/KrakenTools).

To further increase reference database complexity representative of realistic classification scenarios, we utilised an *E. coli* reference database comprising 304 strains (see Supplementary Methods: S2. All-vs-all whole-genome ANI for this 304-strain reference database was computed and visualised using FastANI v1.34[31] (see Data Summary). This informed selection of 10 strains for metagenome spike-in: (i) five equal abundance intra-phylogroup (B2-2) strains of increasing genetic relatedness (ANI), and (ii) five more diverse strains at differential abundances (Table S8). Spike-in reads were generated using InSilicoSeq[30] v2.0.1, corresponding to 1% total paired reads in each metagenome, as determined using SeqKit[32] v2.13.0. Generated spike-in reads were concatenated with their respective metagenomes. Custom scripts for spike-in read generation are available (see Data Summary).

The final 42 spiked metagenomes were used as input for all tools, alongside the 304 *E. coli* reference strain database. To further assess the impact of not including the strains present in the metagenome in the reference database, analyses were repeated using a reference database excluding the 10 *E. coli* strains present in Table S8, leaving 294 strains.

### Computational benchmarking

To compare computational requirements of the six tools under realistic conditions, a computational benchmark was performed using the real-world spike-ins dataset. The */usr/bin/time -v* utility recorded runtime and computational resource usage for each tools database-building step and profiling step for the 42 metagenome samples. Computational benchmarking was performed on a dedicated compute node (2×Intel(R) Xeon(R) E5-2640 v4 @2.40 GHz; 20 physical cores; 40 logical CPUs), with each tool restricted to four threads, selected to represent a resource allocation accessible to users without high-performance computing infrastructure, and memory usage uncapped (503 GiB free). An exception was PanTax profiling, which could not complete using four threads and instead ran on dedicated SLURM-scheduled compute nodes (Intel(R) Xeon(R) Gold 6126 @2.60 GHz), with 32 threads and 400GB memory allocated per job. Runtime and peak memory usage (maximum resident set size) were used for comparative computational benchmarking, with all recorded statistics made available (see Data Summary).

### Performance metrics

Across datasets, estimation metrics were calculated from predicted relative abundances using error, absolute error, absolute proportional error, mean error, mean absolute error and mean absolute proportional error, as appropriate. Detection metrics of sensitivity, specificity, precision and F1 score were calculated based on raw call-type counts (true positives, false positives, false negatives, and true negatives) for the simulated and real-world metagenomes multi-strain spike in datasets only, as the expanded reference genome databases used in these experiments allowed for false positive and true negative strain detections. However, due to false negative detections in the SRR13355226 mock community dataset, sensitivity was calculated. Full equations are available (Supplementary Methods S3). Detection thresholds were applied at relative abundance cut-offs of 0.0001, 0.0005, 0.001, and 0.005. Predicted relative abundances below each threshold were set to zero, and presence/absence calls were redefined for recalculation of sensitivity, specificity, precision, and F1 score (see Data Summary).

### Statistical analysis and visualisations

Statistical analysis and visualisations were performed using R[33] v4.5.1. R packages version information is available (Supplementary Methods: S4). For the SRR13355226 mock community dataset, differences in prediction error between human read removal conditions were assessed using paired Wilcoxon signed-rank tests on absolute error, performed per tool to account for the paired structure of observations (significance threshold: *p*<0.05). For StrainScan within the real-world spike-ins dataset, differences in paired metagenome size between samples in which one versus multiple strains were detected were assessed using a Wilcoxon rank-sum test (significance threshold: *p*<0.05).

## 7. Results

### Strain-level profiling error varies across tools despite equal abundance of five ***E. coli* strains**

Using the ZymoBIOMICS® D6331 gut microbiome standard (low-complexity microbiome, five equal relative abundance *E. coli* strains (Table S2) also included in the reference genome database), all tools predicted differing estimates of relative abundance per strain, except for identical predictions of B2207 from StrainScan and Strainify (0.165; Fig.1a). For 5/6 tools, B1109 and B3008 were consistently overestimated (error range: 0.009–0.034 and 0.001–0.094, respectively; Fig.S1a), while B766 was consistently underestimated (error range: −0.008 to −0.088; Fig.S1a). The exception was PanTax, which predicted identical relative abundances (0.2) for all strains (Fig.1a), resulting in zero error (Fig.S1a). Consequently, PanTax achieved the lowest mean absolute error across strains (0; Fig.1b), outperforming all other tools. StrainGE exhibited the highest mean absolute error (0.065). PathoScope, StrainScan, Strainify and StrainR2, and StrainScan showed intermediate mean absolute errors of 0.006, 0.018, 0.02 and 0.022, respectively (Fig.1b).

### Absolute proportional error in relative abundance predictions increases with decreasing sequencing depth

Using the SRR13355226 mock community (99% human, 1% *E. coli* reads), four *E. coli* strains at differential relative abundances, also included in the reference database) we assessed tool performance at different sequencing depths (Table S3), and the impact of human read removal.

Human read removal drastically reduced file size (1.8/1.9 Gb versus 32/33 Mb for paired-end SRR13355226 and SRR13355226_depleted files, respectively) but had negligible impact on strain-level abundance accuracy (paired Wilcoxon signed rank on absolute error *p*>0.05; Fig.S2a).

All tools except StrainScan detected all four strains across conditions (Fig.2a), achieving perfect sensitivity (Fig.S2b). StrainScan failed to detect O157:H7 str. Sakai and O139:H28 str. E24377A (sensitivity: 0.50; Fig.S2b). StrainScan run in super-low-depth mode detected all strains, although the predicted relative abundances deviated substantially from the ground truth (Fig.2a), indicating a trade-off between sensitivity and accuracy. Whilst all remaining tools preserved the correct ranking of strains by relative abundance, all predictions deviated from the ground truth relative abundance (Fig.2a). An exception was PanTax, which assigned identical relative abundances (0.102) to the two lowest-abundance strains (Fig.2a).

Estimation accuracy was assessed using absolute proportional error (APE) to compare across strains with differing ground truth abundances. Across tools, APE increased as sequencing depth decreased, with exceptions including StrainGE for UTI89 (0.05) and PathoScope for O157:H7 str. Sakai (0.46; Fig.S2c). For detected strains, APE ranged across tools from 0.19–0.48 (H10407) 0.05–1.38 (UTI89), 0.46– 2.99 (O157:H7 str. Sakai), and 14.92–164.15 (O139:H28 str. E24377A) (Fig.S2c). For O139:H28 E24377A, the lowest abundance strain, APE was subjectively influenced by both missed detections (APE=1; StrainScan) and large overestimations by other tools which were heavily penalised. This drove large variation per tool when taking mean APE (Fig.2b). Across strains, StrainScan had the lowest mean APE of 0.89 (Fig. 2b), driven by reduced sensitivity (Fig. S2b). Among the remaining tools, PathoScope was the most accurate (4.08), whilst StrainScan super-low-depth provided the least accurate relative abundance predictions (42.02).

**Fig. 1.**
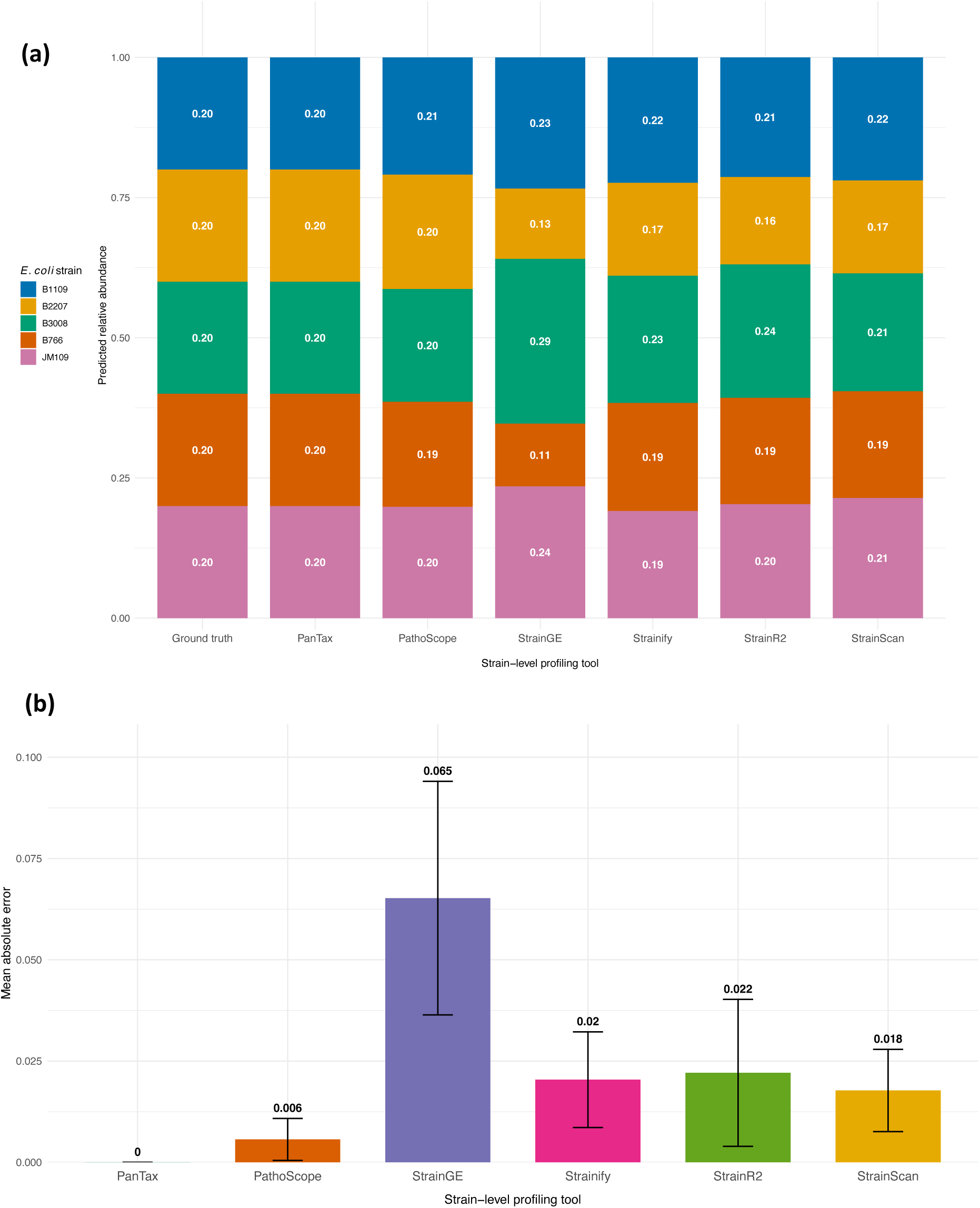
Predicted relative abundance and mean absolute error per strain-level profiling tool for five equally abundant *E. coli* strains in the ZymoBIOMICS® D6331 gut microbiome standard. (a) Predicted relative abundances are shown as stacked bars for each tool. The ground truth relative abundance (0.2 for all strains) is shown on the far left bar. (b) Mean absolute error across all strains (*n* = 5), with error bars +/-standard deviation.

**Fig. 2.**
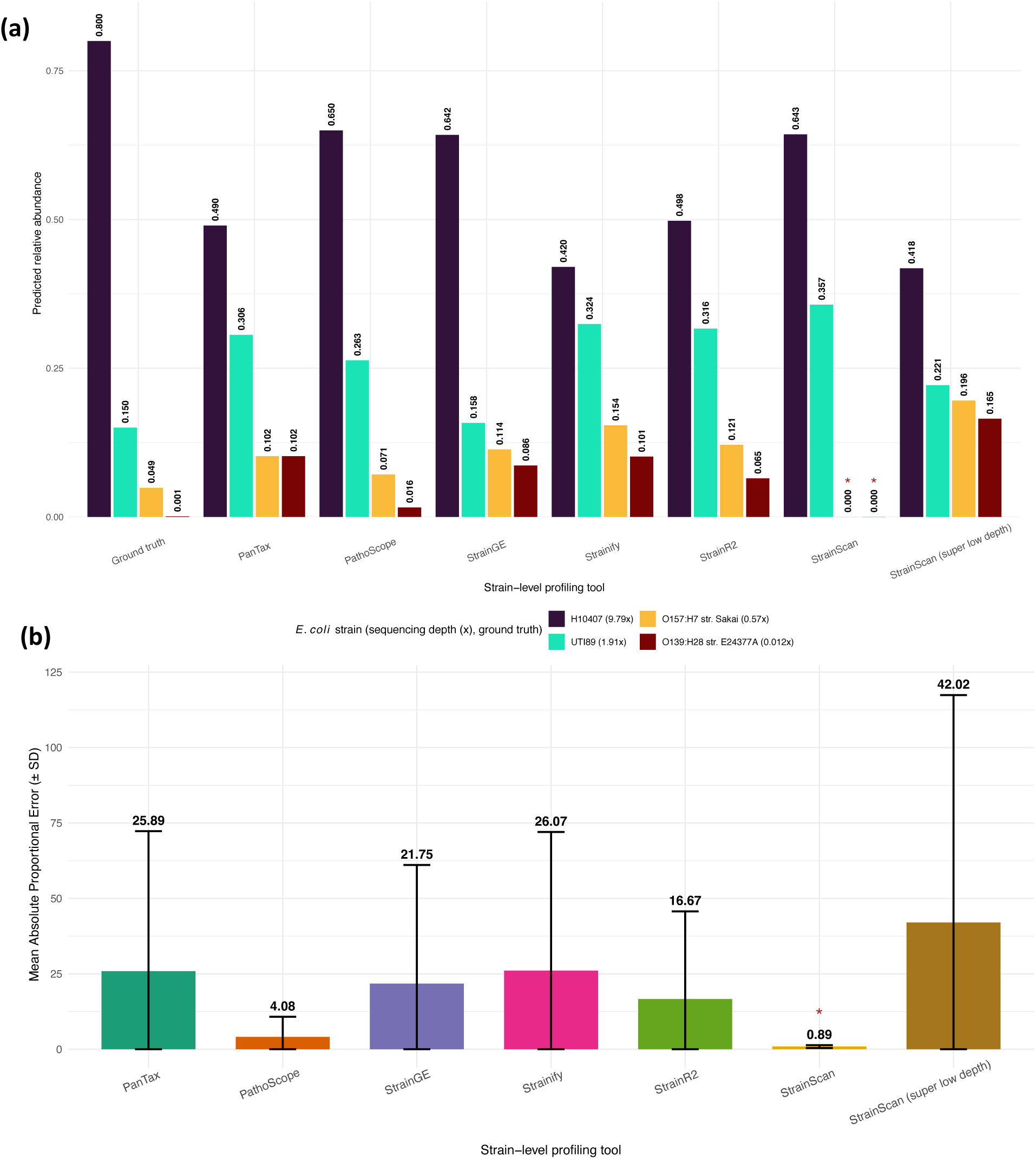
Predicted relative abundance and mean absolute proportional error of four differentially abundant *E. coli* strains in the SRR13355226 mock community across strain-level profiling tools. All tools were run on datasets with (SRR13355226_depleted) and without (SRR13355226) human read removal and collapsed by base tool as no significant differences were observed between human read removal conditions (Fig.S2a). (a) Raw predicted relative abundances by tool. Values are shown relative to the ground truth (far left). An asterisk (*) indicates that a strain was not detected by the corresponding tool. (b) Mean absolute proportional error (± SD) per base tool across all four strains and both conditions (*n* = 8). An asterisk (*) indicates that StrainScan failed to detect O157:H7 str. Sakai (0.57×) and O139:H28 str. E24377A (0.012×) across conditions; these observations were assigned an APE of 1 (see Methods).

### Detection thresholds selectively improve strain-level detection in simulated gut metagenomes

To increase benchmarking complexity, we simulated metagenomes reflecting a baseline healthy adult gut microbiome with two *E. coli* strains (K12-MG1655 and O157:H7 Sakai) spiked across a range of differential abundances and sequencing efforts (Table S4). We also increased the complexity of the reference database to 24 *E. coli* strains representing diversity within and across major phylogroups (Table S5).

Summary detection metrics of 54 simulated metagenomes showed that strains that were genuinely present (true positives) were rarely, if ever, missed, with all tools exhibiting high sensitivity ranging from 0.833 (StrainScan) to 1.000 (PathoScope, Strainify, and StrainR2) (Fig.3a; Table S8). This was true even at low sequencing depth of O157:H7 str. Sakai (≥0.210×; Table S4). However, the reported detection of absent strains (false positives) was a problem for some tools, with PathoScope (0.014; 0.070), Strainify (0.054; 0.073), and StrainR2 (0.596; 0.160) exhibiting lower specificity and precision (Fig.3a; Table S8). Other tools exhibited high specificity and precision, including StrainGE (0.998; 0.967), StrainScan (1.000; 1.000), and StrainScan super-low-depth mode (1.000; 1.000) (Fig.3a; Table S8).

The F1 score, defined as the harmonic mean of sensitivity and precision, provides a balanced summary of detection performance. StrainGE achieved the highest F1 score (0.978), followed by StrainScan super-low-depth (0.947), StrainScan (0.909) and PanTax (0.816) (Fig.3a; Table S8). The remaining tools showed lower F1 scores (StrainR2, 0.270; Strainify, 0.136; PathoScope, 0.131).

Applying increasing detection thresholds (0.0001–0.005) led to increases in specificity, precision, and F1 score for PathoScope, Strainify, and StrainR2 (Fig.3b). F1 scores increased to 0.733 (PathoScope), 0.466 (Strainify), and 0.952 (StrainR2) at the 0.005 detection threshold, with the latter exceeding both StrainScan modes original F1 score (Fig.3b). As sensitivity remained unchanged, these increases were driven by removal of numerous low-abundance false positive detections from the 24-strain *E. coli* reference database. In contrast, PanTax, StrainGE, StrainScan, and StrainScan super-low-depth mode were unaffected by the applied detection thresholds (Fig.3b).

### Predicted relative abundances in a two-strain simulated gut metagenome scenario are generally accurate

We compared predicted relative abundances for K12-MG1655 and O157:H7 str. Sakai across simulated gut metagenomes (Fig.4). PathoScope and Strainify reported predicted relative abundance for the negative control (both strains at zero abundance), with assignments predominantly to the “Other” strain category, representative of 22 false-positive *E. coli* strains within the reference database (Fig. 4a; Table S5). However, these alignment-based tools exhibited lower proportions of mapped reads in the negative control compared to all other samples: PathoScope assigned 0.04% of reads in the negative control, compared to 1.02% across the remaining samples, while Strainify assigned 0.02% versus 1.60–1.77%, respectively (Fig.S3a). Other tools did not report predicted relative abundance for the negative control (Fig.4a).

**Fig. 3.**
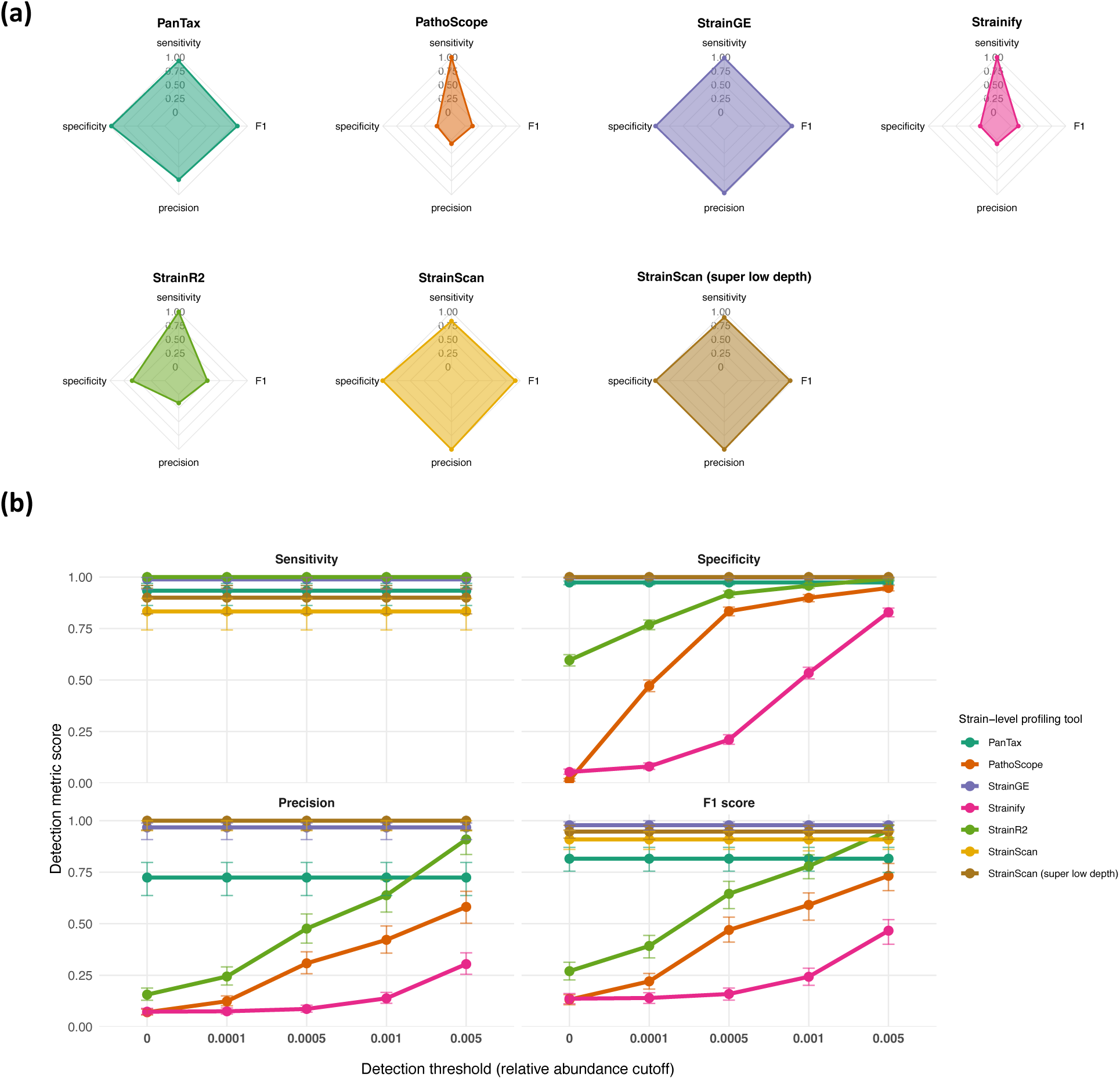
Detection metrics (sensitivity, specificity, precision, and F1 score) per tool are differentially affected by increasing detection thresholds when using simulated metagenomes. (a) Detection metrics summarised as radar plots for each tool across all simulated metagenomes (*n =* 54), with call-type counts pooled across all metagenomes to produce a single aggregate value for each metric per tool. Metrics were computed from raw true positive, false positive, true negative, and false negative call-type counts per tool (*n* = 1,296; see Data Summary). All metrics are scaled between 0–1. b) Detection metric scores per tool across increasing detection thresholds in the simulated metagenomes dataset. Thresholds (0.0001–0.005; discrete *x* axis shown) were applied to raw call-type counts for all tools, and call types were reassigned accordingly (see Data Summary) and re-pooled across all metagenomes to produce a single aggregate value for each metric per tool. Error bars indicate 95% confidence intervals (Wilson intervals for sensitivity, specificity, and precision; bootstrap intervals for F1 scores).

**Fig. 4.**
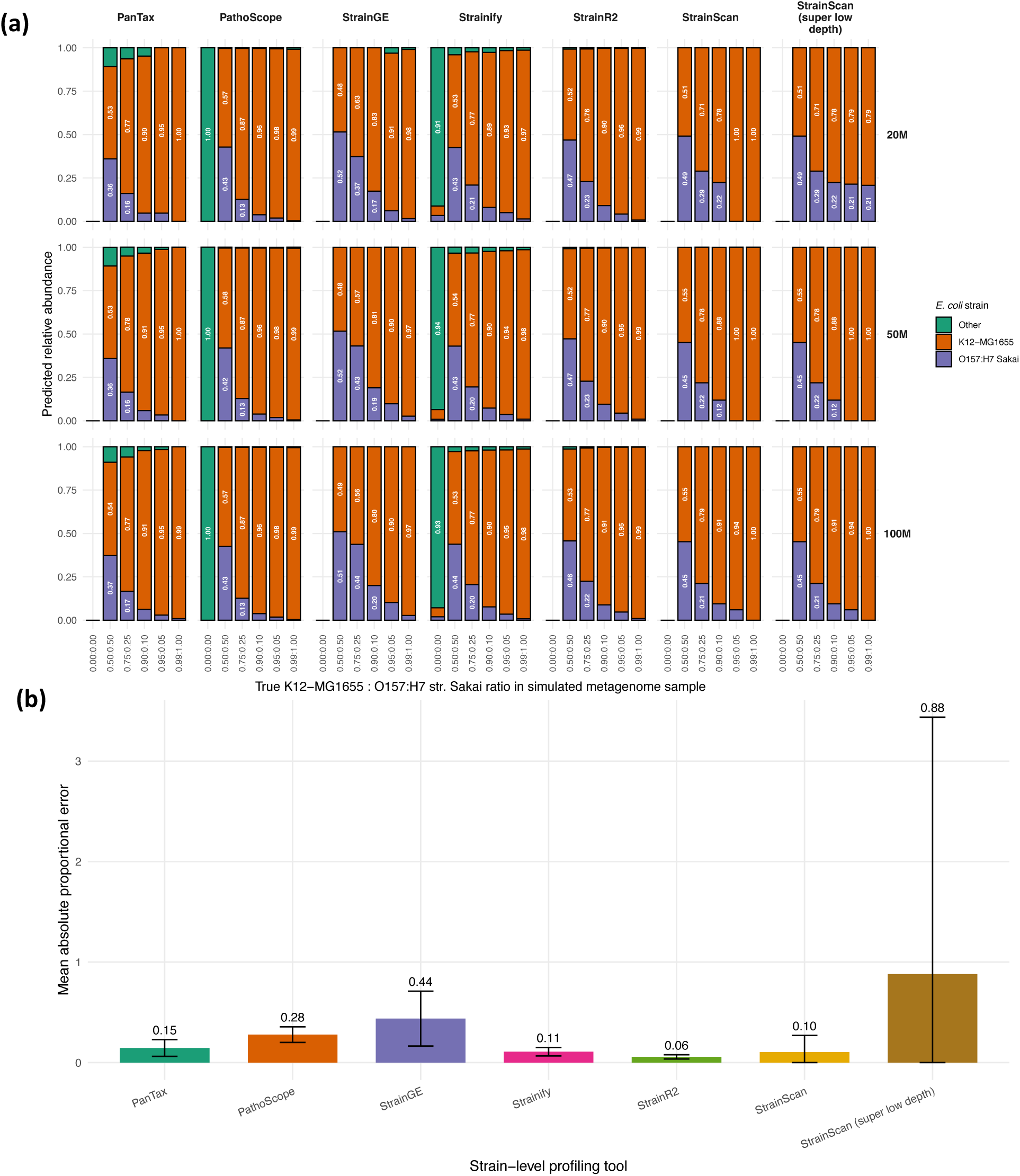
Predicted relative abundance and mean absolute proportional error per tool for the simulated metagenome dataset. (a) Predicted relative abundance per simulated metagenome sample, stratified by tool and sequencing effort. The true ratio of *E. coli* K-12 MG1655 (orange) to O157:H7 str. Sakai (purple) is shown on the x-axis, with predicted relative abundances displayed as stacked bars on the y-axis. “Other” (green) denotes predicted relative abundance assigned to the remaining 22 false-positive *E. coli* strains in the reference database (Table S5). The 0.00:0.00 sample represents a negative control. All values represent means across triplicates (see Methods), and no detection thresholds were applied. (b) Mean absolute proportional error per tool, calculated from the predicted relative abundances of K-12 MG1655 and O157:H7 str. Sakai. Error bars represent mean ± standard deviation.

Across the remaining metagenome samples, PanTax and Strainify showed the highest predicted relative abundances assigned to false-positive “Other” strains (PanTax: 0.014–0.120, *n* = 32; Strainify: 0.000039–0.0930, *n* = 1,141; Fig.4a; see Data Summary). For PanTax, these fewer, higher-abundance false-positive assignments were solely assigned to O157:H7 str. EDL933, the closest reference to O157:H7 str. Sakai by ANI (99.97%; Table S6). Consequently, all PanTax false-positive assignments exceeded the evaluated detection thresholds (0.0001–0.005), explaining why increasing thresholds had no effect on its detection metrics (Fig. 3b).

All tools demonstrated strong performance across sequencing efforts, with predicted relative abundances closely reflecting the expected stepwise increase in abundance of K12-MG1655 (orange) and decrease in O157:H7 str. Sakai (purple). StrainR2 showed the closest agreement with the expected ratios (Fig.4a). Exceptions included PanTax at 20M and 50M (0.99:0.01), and StrainScan at 20M and 50M (0.95:0.05, 0.99:0.01), and 100M (0.99:0.01), which all missed detection of O157:H7 str. Sakai at these lower sequencing depths (0.21–2.63x; Table S4). StrainScan low depth mode recovered detection of O157:H7 str. Sakai in the 20M 0.95:0.05 and 0.99:0.01 samples (Fig.4a).

Taking mean APE of the ground truth strains across all simulated metagenomes, StrainR2 was the most accurate tool (0.06; Fig.4b). StrainScan (0.10), Strainify (0.11), PanTax (0.15), PathoScope (0.28), and StrainGE (0.44) showed intermediate errors. StrainScan super-low-depth mode had the highest mean APE (0.88), driven by rescued detection of O157:H7 str. Sakai in the 20M 0.95:0.05 and 0.99:0.01 samples, where predicted abundances (0.79:0.21; Fig.4a) deviated strongly from the ground truth. Across all tools, when splitting mean APE by strain, K12-MG1655 showed consistently lower mean APE, ranging from 0.02 (StrainR2) to 0.08 (StrainGE), whereas O157:H7 Sakai showed consistently higher mean APE, ranging from 0.10 (StrainR2) to 2.11 (StrainScan super-low-depth) (Fig.S3b), indicating that overall error was driven by the strain with lower sequencing depth (Table S4).

### Removing true strains from the reference database leads to out-of-phylogroup assignments for some tools when using simulated metagenomes

In some ‘real-life’ applications, the exact strain present in a metagenomic sample will be absent from the reference database. To simulate this, we removed K12-MG1655 and O157:H7 Sakai (phylogroups A and E; Table S5) genomes from the database, leaving 22 ‘false-positive’ *E. coli* strains. All tools were rerun on the same simulated gut metagenomes to assess how predicted relative abundances were reassigned (Fig.5).

**Fig. 5.**
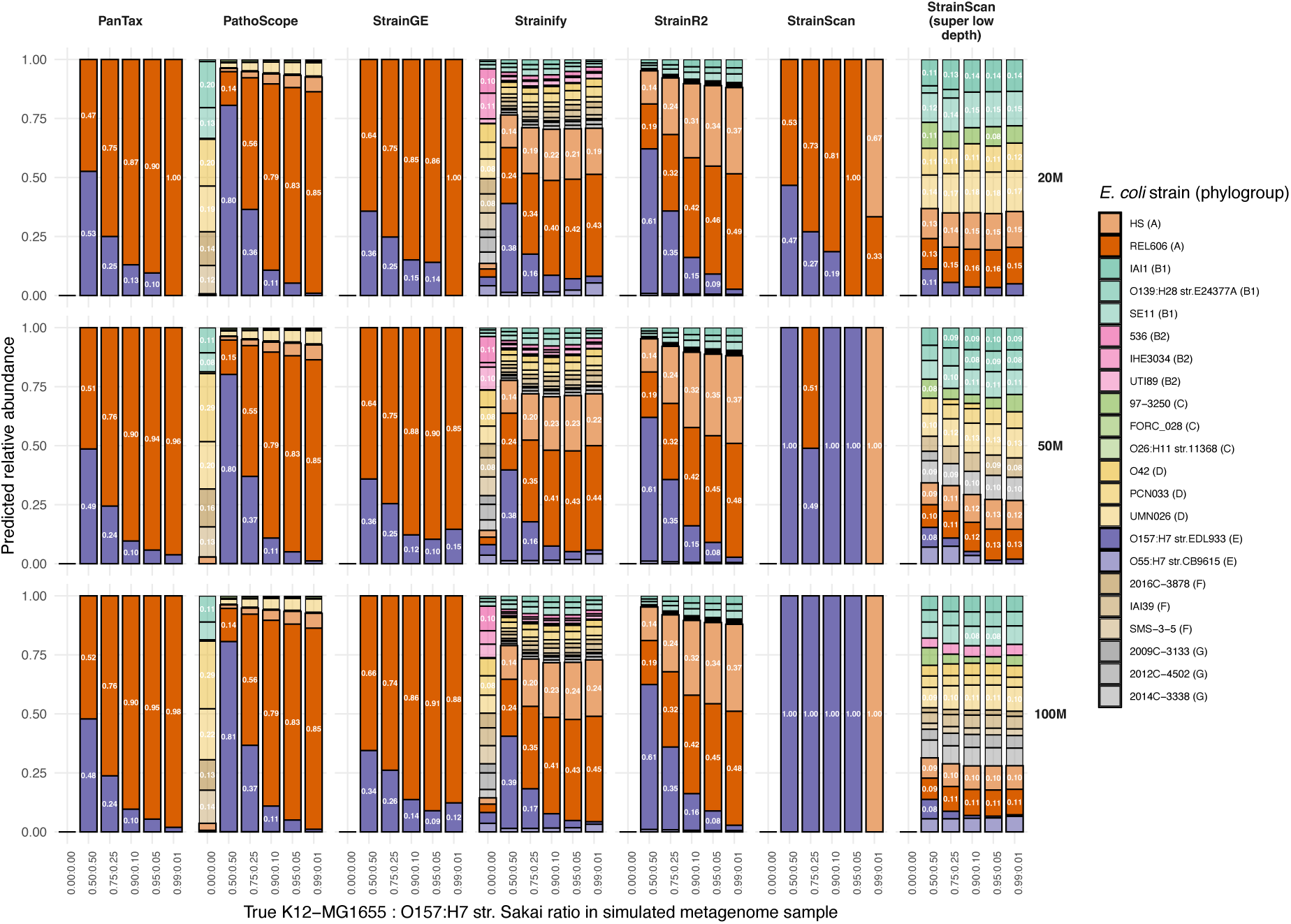
Predicted relative abundance per tool across simulated metagenomes when K12-MG1655 and O157:H7 str. Sakai are excluded from the reference database. Predicted relative abundances are shown for each simulated metagenome, faceted by tool and sequencing effort. The two true strains (K12-MG1655 and O157:H7 str. Sakai) were removed from the reference database prior to analysis, while the remaining 22 false-positive strains were unchanged (Table S5). The initial ground truth K12-MG1655 (phylogroup A) to O157:H7 str. Sakai (phylogroup E) ratio is shown on the x-axis; the 0.00:0.00 sample represents a negative control. Stacked bars indicate predicted relative abundances, with phylogroups A and E coloured as in previous figures (orange and purple, respectively; Fig.4a). Values represent means across triplicates (see Methods). No detection thresholds were applied.

PanTax most closely mirrored the expected phylogroup-level proportions, missing a phylogroup E assignment (O157:H7 str. EDL933/O55:H7 str. CB9615) only in the 20M 0.99:0.01 sample. In all other samples, phylogroup A and E predictions were consistently assigned to REL606 and O157:H7 str. EDL933, respectively, which were the closest available reference strains to K12-MG1655 (98.95% ANI) and O157:H7 Sakai (99.97% ANI) (Table S6). StrainGE also closely preserved the expected phylogroup A:E abundance ratios, with stepwise increased assignment to REL606 and corresponding decreases in O157:H7 str. EDL933, except at 0.99:0.01. PathoScope showed similar phylogroup-level accuracy but split phylogroup A abundance between REL606 and HS (98.74% ANI), with the remaining abundance (≤0.07) assigned predominantly to phylogroup D strains and numerous low-abundance strains.

StrainR2 broadly recapitulated these phylogroup-level patterns, but distributed phylogroup A abundance more evenly between REL606 and HS, with increasing assignment to B1 strains (≤0.10) as the true K12-MG1655 ratio increased. Strainify showed a similar trend but redistributed ≤0.35 predicted relative abundance across a broader range of phylogroups (B1, B2, C, D). Again, only Strainify and PathoScope produced relative abundance predictions for the negative control.

At 20M depth, StrainScan matched the expected phylogroup ratios for the 0.50:0.50– 0.90:0.10 samples. At 0.95:0.05, all relative abundance was assigned to REL606 (1.00) and at 0.99:0.01 it was split between HS and REL606 (0.67:0.33; Fig.5). At higher sequencing efforts (50M; 100M), predictions were dominated by O157:H7 str. EDL933. In contrast, StrainScan low-depth mode did not reflect the expected phylogroup-level abundance ratios. Relative abundance predictions were distributed broadly across a range of phylogroups (A, D C and B1), with no individual strain exceeding 0.18 relative abundance.

### Detection thresholds remain selectively beneficial across tools for strain-level detection in real-world metagenomes

Whilst simulated gut metagenomes are useful for benchmarking, they may not capture the complexity of “real-life” gut metagenomes. Therefore, we generated the real-world spike-ins dataset, consisting of 42 publicly available short-read gut metagenomes (Table S7) of variable file sizes (Fig.S4). These were depleted of *E. coli* reads and spiked at 1% total reads with 10 *E. coli* strains of varying ANI relatedness with differential abundance (Table S8). To better represent realistic classification scenarios, we expanded the reference genome database to 304 *E. coli* strains.

Analysis of raw detection metrics across the 42 metagenome samples (Fig.6a) revealed greater variation in sensitivity between tools than observed for the simulated metagenomes. PathoScope, Strainify, StrainR2, StrainScan (super-low-depth) and PanTax maintained high sensitivity (1.000, 0.988, 1.000, 0.933 and 0.862 respectively), whereas StrainGE (0.500) and StrainScan (0.424) showed lower sensitivity, frequently failing to detect all 10 true-positive strains. Conversely, StrainGE, StrainScan and PanTax achieved perfect specificity and precision (both 1.000, per tool), while StrainR2 (0.987; 0.717) and StrainScan super-low-depth mode (1.000; 0.987) also performed strongly, indicating none or few false-positive detections and high confidence in positive prediction calls from the 304-strain reference database. PathoScope and Strainify retained low specificity (0.282; 0.687) and precision (0.0453; 0.0869). Overall, StrainScan super-low-depth mode achieved the highest F1 score (0.960), followed by PanTax (0.926; Fig 6a).

**Fig. 6.**
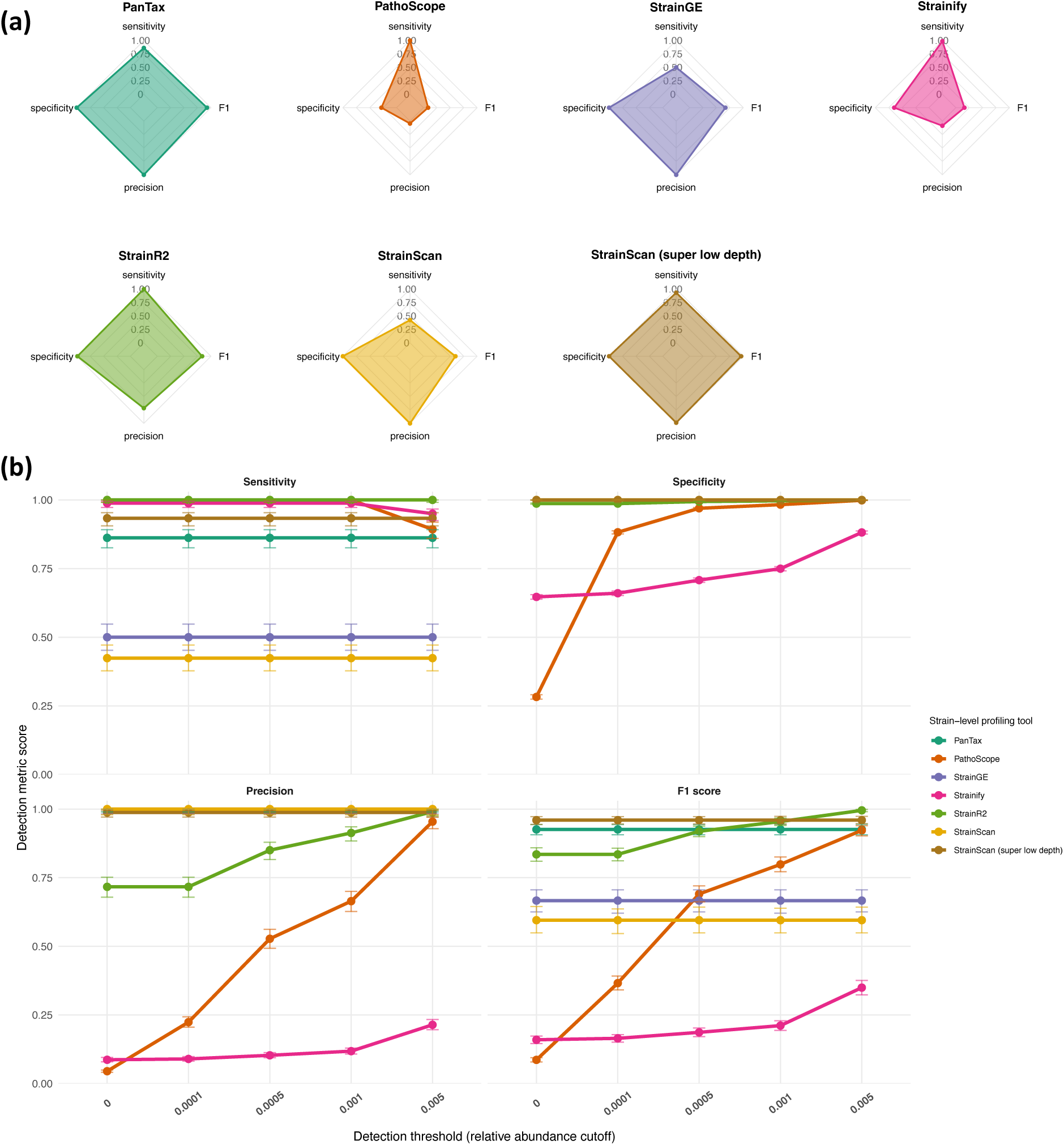
Detection metrics (sensitivity, specificity, precision, and F1 score) per tool are differentially affected by increasing detection thresholds in real metagenomes spiked with 10 *E. coli* strains. (a) Detection metrics summarised as radar plots for each tool in the real-world spike-ins dataset, with call-type counts pooled across all metagenomes (*n* = 42) to produce a single aggregate value for each metric per tool. Metrics were computed from raw true positive, false positive, true negative, and false negative call-type counts per tool (*n* = 12,768; see Data Summary). All metrics are scaled between 0–1. (b) Detection metric scores per tool across increasing detection thresholds in the real-world spike-ins dataset. Thresholds (0.0001–0.005; discrete *x* axis shown) were applied to raw call-type counts for all tools, and call types were reassigned accordingly (see Data Summary) and re-pooled across all metagenomes to produce a single aggregate value for each metric per tool. Error bars indicate 95% confidence intervals (Wilson intervals for sensitivity, specificity, and precision; bootstrap intervals for F1 scores).

Applying detection thresholds (0.0001–0.005) affected only PathoScope, Strainify and StrainR2 (Fig. 6b), consistent with observations from the simulated gut metagenomes dataset. Specificity, precision and F1 score increased with detection threshold, with F1 scores reaching 0.923, 0.350 and 0.995, respectively, at the 0.005 cutoff. StrainR2 therefore achieved the highest F1 score observed in the experiment, mirroring its performance in simulated metagenomes. At this threshold, sensitivity decreased for PathoScope (0.893) and Strainify (0.950), indicating some true-positive predictions fell below the threshold. The remaining tools were unaffected by changing detection thresholds.

### Predicted relative abundances in multi-strain spiked gut metagenomes show variation in accuracy across tools

We compared predicted relative abundances for 10 spiked *E. coli* strains across 42 real-world gut metagenomes (Fig.7). In contrast to the two-strain simulated gut metagenomes, relative abundance predictions varied substantially between tools (Fig.7a). StrainR2 most consistently reproduced the expected strain proportions across samples, with only a notable false-positive assignment to “Other” strains in SRS12879160 (0.042 across six strains). As all 10 strains were consistently detected, StrainR2 resolved strains separated by up to 99.9% ANI (PMV-1 and IHE3034; Fig.S5) across diverse metagenomic backgrounds.

**Fig. 7.**
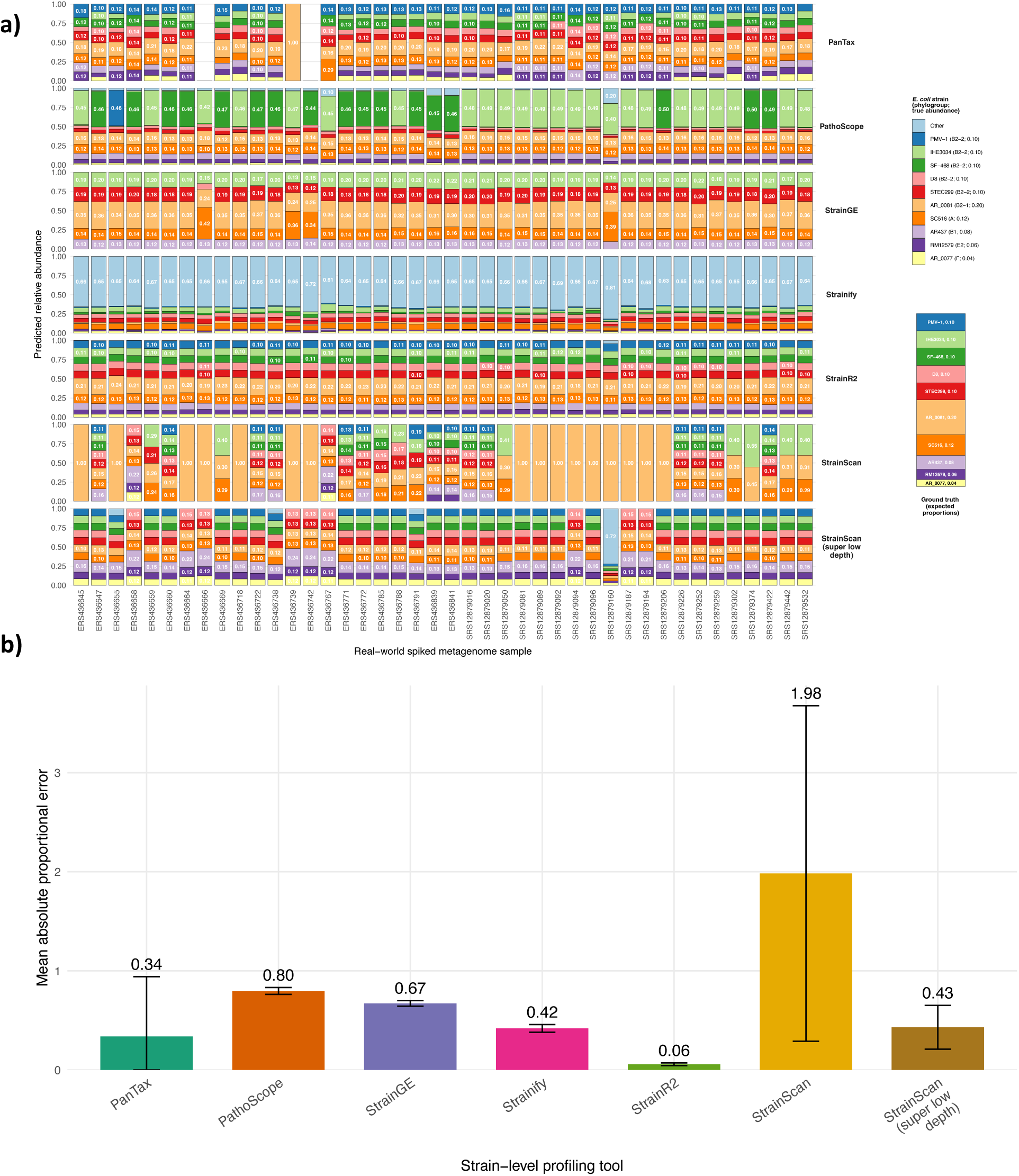
Predicted relative abundance and mean absolute proportional error per tool for the real-world spike-ins metagenomic dataset. (a) Predicted relative abundance per spiked metagenome sample, faceted by tool. Ground-truth relative abundances for the 10 spiked *E. coli* strains are shown alongside an expected ground truth, stacked-bar profile. If predicted relative abundance calls were ≥0.10, assignments were labelled accordingly. “Other” (light blue) denotes the predicted relative abundance assigned to any of the remaining 294 false-positive *E. coli* strains in the reference database (see Data Summary). No detection thresholds were applied. (b) Mean absolute proportional error per tool, calculated from the predicted relative abundances of the 10 spiked *E. coli* strains (Table S9). Error bars represent mean ± standard deviation.

PanTax, PathoScope, Strainify and StrainScan (super-low-depth) also showed high sensitivity (Fig.6a) and so also frequently resolved the most closely related spiked strains. However, their abundance estimates were less accurate than StrainR2 (Fig.7a). PanTax detected all 10 spiked strains in 24/42 samples, with predicted relative abundances closely matching the expected proportions. Of the remaining samples, 15/42 missed at least one of the three lowest-abundance strains (AR437, 0.08; RM12579, 0.06; AR_0077, 0.04), ERS436749 detected only the dominant AR_0081 strain (0.20), and ERS436666 and ERS436742 produced no relative abundance predictions. PathoScope consistently overestimated either IHE3034 or SF-468 (expected: 0.10, predicted: 0.42–0.50), while Strainify predictions were dominated by false-positive “Other” strains (0.63–0.81). StrainScan super-low-depth mode generally reproduced the expected proportions more closely, although nine samples contained only six of the 10 spiked strains, and SRS12879160 showed a large false-positive “Other” assignment (0.72).

StrainScan performed substantially worse, with 16/42 samples entirely assigning predicted abundance (1.00) to the dominant B2-1 strain AR_0081 (0.20; Fig.7a). As StrainScan largely produced these dominant-strain-only profiles, we examined whether detection was associated with metagenome size (Fig.S6). Samples in which only one strain was detected were significantly smaller than those with multiple detections (Wilcoxon rank-sum test, *p* = 6.97 × 10⁻⁶; Fig.S6), indicating reduced strain recovery at lower sequencing depth.

StrainGE showed a different pattern, consistently detecting five of the 10 spiked strains in all 42 samples (Fig.7a). These predominantly comprised IHE3034 and STEC299 from the five closely related B2-2 strains, together with AR_0081 (B2-1), SC516 (A) and AR437 (B1), with D8 detected instead of STEC299 in ERS436666.

These differences were reflected in abundance error (Fig.7b). StrainR2 had the lowest APE (0.06), followed by PanTax (0.34), Strainify (0.42), StrainScan super-low-depth mode (0.43), StrainGE (0.67), PathoScope (0.80) and StrainScan (1.98).

### Tools differentially retain phylogroup-level information in real-world metagenomes when true strains are removed from the reference database

To determine how predictions were reassigned at the phylogroup level when exact strain references were unavailable, the 10 spiked ground-truth *E. coli* strains (Table S8) were removed from the reference database, leaving 294 “false positive” strains, and all tools rerun across the same 42 metagenomes (Fig.8). To minimise low-abundance false positives associated with the denser reference database, we applied a 0.005 relative abundance detection threshold.

**Fig. 8.**
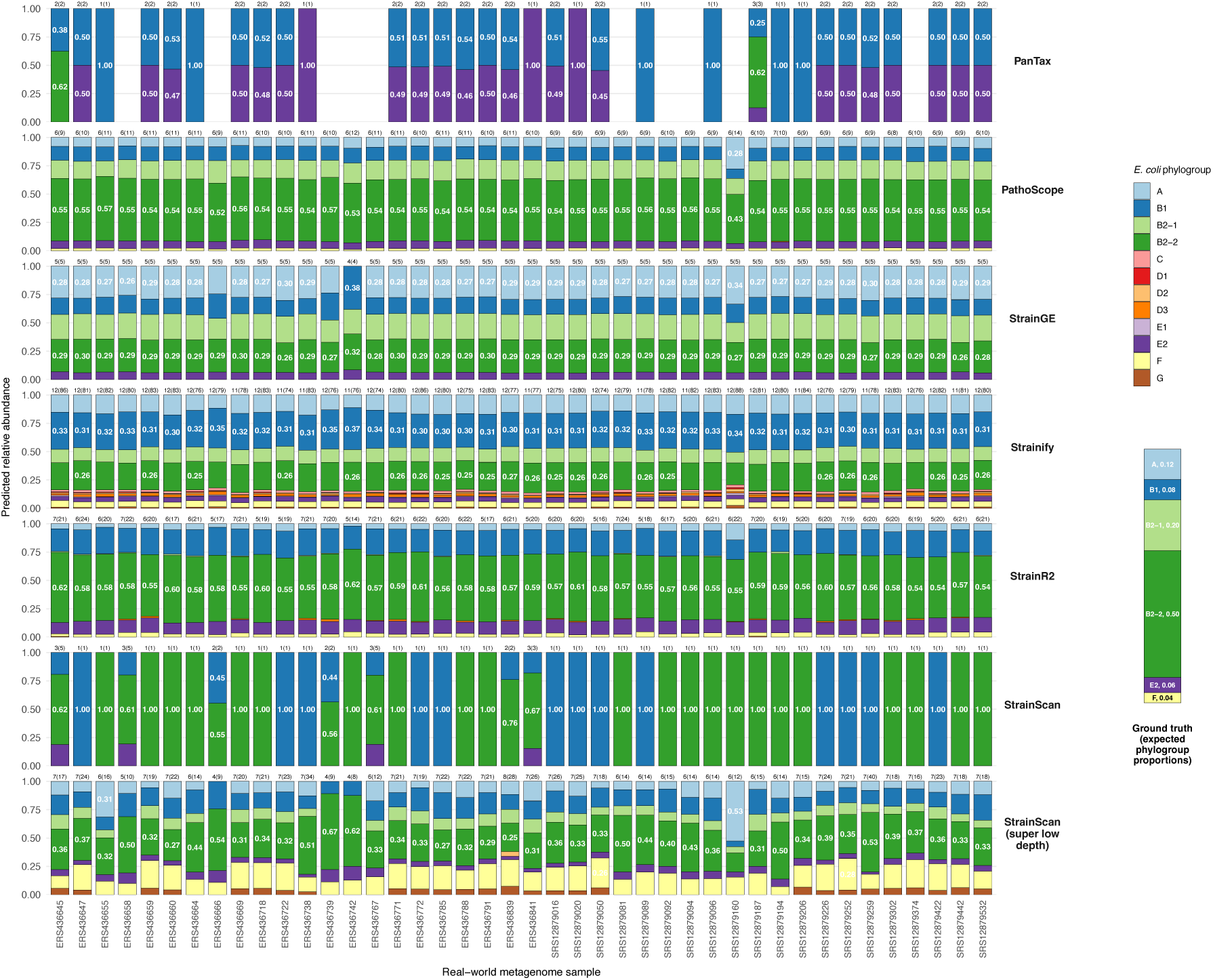
Phylogroup-level predicted relative abundance per tool across spiked metagenomes when ground truth strains are excluded from the reference database. Predicted relative abundances at the phylogroup level are shown for each metagenome (*n* = 42) in the real-world spike-ins dataset, faceted by tool. The 10 spiked ground-truth strains (Table S9) were removed from the reference database prior to analysis, leaving 294 strains in the reference database (see Data Summary). The expected ground truth phylogroup relative abundance stacked-bar profile is shown. Bars were labelled if predicted relative abundance assignments were ≥0.25. Numbers above individual stacked bars indicate the number of phylogroups detected (number of strains detected). A detection threshold of 0.005 was applied to remove numerous low abundance “false positive” detections. Values were renormalised after application of the detection threshold (see Data Summary).

At this threshold, PathoScope showed the strongest agreement with the expected phylogroup composition. In 41/42 samples, PathoScope detected exclusively the six expected ground-truth phylogroups; the exception, SRS12879160, contained one additional phylogroup. PathoScope reported 8–12 individual strains per sample, which was substantially fewer than the 136–273 strains spanning all 12 phylogroups detected before thresholding (see Data Summary). StrainGE also performed strongly but was more conservative, consistently recovering five of the six expected phylogroups in 41/42 samples. Its predicted proportions showed less agreement with the expected composition, and the lowest abundance phylogroup, F (expected: 0.04), was consistently absent. Across all samples, however, StrainGE reported only a single strain per detected phylogroup.

Amongst other tools, StrainR2 recovered the dominant B2-2 phylogroup across all samples (expected: 0.50; predicted: 0.54–0.62), but consistently underestimated B2-1, while detecting 5–7 phylogroups and 16–24 individual strains per sample. StrainScan predominantly produced single-strain, dominant-phylogroup predictions, alternating mainly between B2-2 and B1; only seven samples contained multiple predicted phylogroups. StrainScan super-low-depth mode recovered substantially greater phylogroup diversity, although predictions varied across samples, with B2-2 remaining dominant in 41/42 samples (0.25–0.67). Strainify detected all 12 possible phylogroups and a high number of individual strains (74–86), whilst incorrectly assigned B1 as dominant (expected: 0.08; predicted: 0.30–0.37). PanTax produced no predictions for 10/42 samples. Predictions for the remaining 32 samples comprised up to three strains/phylogroups: APEC O18 (B2-2), 2013C-4465 (E2), and FDAARGOS_443 (B1), with the latter two typically accounting for approximately equal proportions (∼50:50).

As PathoScope and StrainGE showed the strongest performance, we further examined the genomic relatedness of their predictions to the 10 removed ground-truth strains (Table S9). For each ground-truth strain, we identified the detected reference with the highest pairwise ANI across all samples. PathoScope recovered the closest available reference for eight of 10 strains (99.05–99.90%); the exceptions were STEC299 and AR_0077, whose closest predictions ranked fifth (98.86%) and sixth (98.47%), respectively. StrainGE recovered the closest reference for six of 10 strains.

Its B2-1 prediction, O25b:H4, ranked fourth (99.83%), while the five B2-2 strains collapsed to a single prediction, APEC O18, which ranked first for three strains (99.09– 99.90% ANI) and fourth or fifth for SF-468 (99.67%) and STEC299 (98.86%), respectively. No phylogroup F strain was recovered. Despite identifying closer references overall, PathoScope predictions were more variable across samples: the 10 closest predictions were detected in 4–42 metagenomes, with 31 unique strains detected overall (see Data Summary). StrainGE produced only five unique predictions (Table S9), which were consistently recovered across 41/42 metagenomes.

### Most tools have computational resource requirements accessible to users without high-performance computational infrastructure

We recorded the computational resource required whilst building the 304 *E. coli* strain reference database and profiling of the 42 metagenomes from the real-world spike-ins dataset. Wall-clock runtime and peak memory usage (PMU) were reported (Fig.9), with all recorded metrics available (see Data Summary).

**Fig. 9.**
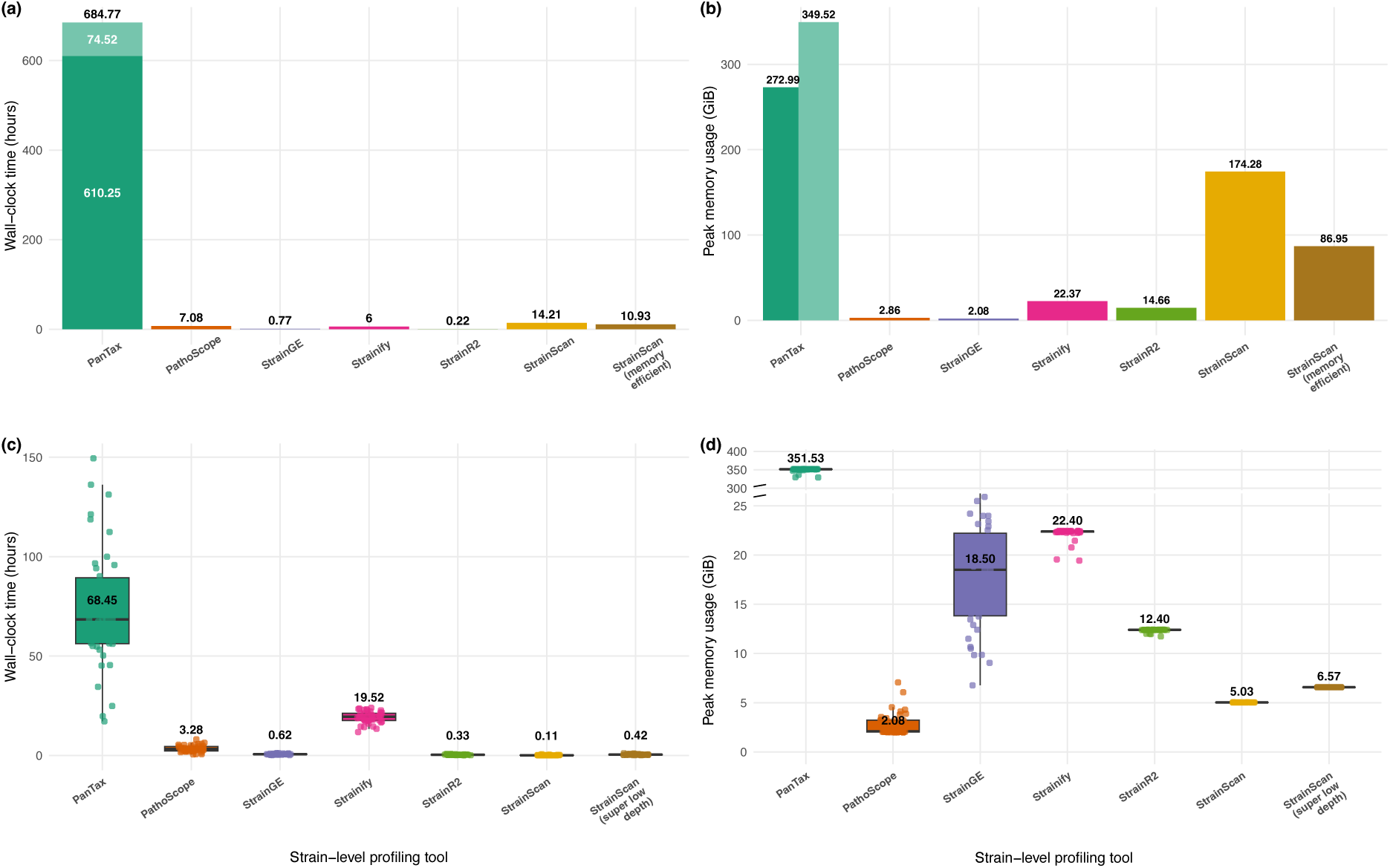
Wall-clock runtime and peak memory usage of tools during database building and profiling of 42 spiked metagenomes. Computational resource usage was recorded using the */usr/bin/time -v* command line utility, with wall-clock runtime (hours) and peak memory usage (maximum resident set size; GiB) shown for database building using the 304-strain *E. coli* reference database (a, b) and profiling of each of the 42 real-world spike-in metagenomes (c, d). Computational resource statistics were recorded using 4 threads, except for PanTax metagenome profiling (c, d), which required 32 threads (see Methods). Database-building measurements (a, b) are shown as bars as each database was built once. Database-building commands used for each tool are provided in the Data Summary. PanTax comprised two independent database-building steps: pangenome graph generation (dark green) and indexing (light green). Their wall-clock times are stacked in (a) to represent the total sequential database-building time, whereas peak memory usage is shown separately in (b) as peak memory usage was measured independently for each step. Profiling measurements (c, d) are shown as box plots with individual metagenome measurements overlaid (*n* = 42, per tool). Values displayed on the box plots (c, d) represent medians. Parallel dashed lines in (d) indicate a *y*-axis break.

For most tools, database building was relatively fast and required modest memory (Fig.9a/b). StrainR2 and StrainGE completed in under one hour (0.22 and 0.77 h, respectively), while Strainify and PathoScope required 6.00 and 7.08 h. PMU across these four tools remained comparatively low, ranging from 2.06 GiB (PathoScope) to 22.37 GiB (Strainify). StrainScan was more resource intensive, requiring 14.21 h and 174.28 GiB in standard mode, reduced to 10.93 h and 86.95 GiB in memory-efficient database-building mode. PanTax had the greatest computational requirements on 4-threads, requiring 684.77 h (28.53 days) to complete database-building, with PMU reaching 349.52 GiB during indexing.

The high computational requirements of PanTax continued during metagenome profiling (Fig.9c/d), with samples unable to complete using four threads. Profiling was instead performed using 32 threads, yet PanTax required a median runtime of 68.45 h and PMU of 351.53 GiB, substantially greater than the other tools. Among the remaining tools, four had median runtimes below one hour: StrainScan (0.11 h), StrainR2 (0.33 h), StrainScan super-low-depth mode (0.42 h) and StrainGE (0.62 h). PathoScope had an intermediate median runtime of 3.28 h, while Strainify required 19.52 h, with both showing greater variability across differentially sized metagenomes (Fig.S4). Median PMU among these tools ranged from 2.08 GiB for PathoScope to 22.40 GiB for Strainify, with StrainScan (5.03 GiB), StrainScan super-low-depth mode (6.57 GiB), StrainR2 (12.40 GiB) and StrainGE (18.50 GiB) falling between these extremes.

StrainR2 was the only tool allowing memory allocation control during profiling, enabling further assessment across allocations of 8–64 GB (Fig.S7). Profiling failed at 8 GB, but median runtime remained similar across successful settings (0.27–0.36 h; Fig.S7a), while PMU decreased with lower memory allocations (Fig.S7b). Thus, StrainR2 could operate across a range of memory allocations with minimal impact on runtime.

## 8. Discussion

We benchmarked six tools (PanTax, PathoScope 2.0, StrainGE, Strainify, StrainR2 and StrainScan) for strain-level profiling *Escherichia coli* within short-read gut metagenomes. We evaluated strain detection and relative abundance estimation across metagenomic datasets of increasing complexity, each designed to test different aspects of strain-level profiling. Across these datasets, tools varied substantially in both the number and relative abundance of strains predicted, including when ground-truth strains were absent from the reference database. Computational requirements also varied considerably between tools, although most could be run with resources accessible without high-performance computing infrastructure.

To our knowledge, this represents the first independent *E. coli*-specific benchmark of short-read metagenomic strain-level profiling tools. The ZymoBIOMICS® D6331 dataset has not previously been used for this purpose, while our simulated gut metagenomes and real-world spike-ins datasets were generated independently. Although the SRR13355226 mock community has been used previously[22, 23, 27], it had not been evaluated across this tool combination. For example, the Strainify authors used its four *E. coli* strains to construct simulated datasets[23], and reported findings consistent with ours: StrainScan struggled to detect low-abundance strains, while super-low-depth mode improved detection but substantially overestimated abundance. However, the impact of human read removal on relative abundance estimation was not assessed. Human read removal had no significant effect on abundance prediction accuracy across tools. Given host DNA typically comprises ∼10% of faecal samples[34], this supports human read removal prior to strain-level profiling without compromising relative abundance estimates, while simultaneously reducing storage requirements and limiting retention of potentially identifiable human sequence data.

As dataset complexity increased, distinct differences in tool behaviour emerged. Across simulated gut metagenomes containing two true strains, all tools exhibited high sensitivity (0.833–1.000). However, when complexity increased to 10 strains within the real-world spike-ins dataset, StrainGE and StrainScan exhibited reductions in sensitivity. Notably, the 10 strains included pairs sharing up to 99.9% ANI, equivalent to approximately one nucleotide difference per 1,000 aligned bases. At this level of genomic similarity, distinguishing strains using short reads becomes inherently challenging: for ∼150-bp metagenomic short-reads, most reads may map equally well to both genomes[35]. Despite this, several tools successfully detected most or all 10 strains, with StrainR2 consistently detecting all 10. Although these experiments do not establish absolute ANI thresholds for strain resolution per tool, they demonstrate that some approaches can distinguish multiple highly related strains.

However, the ability to detect multiple true positive strains must be balanced against specificity and precision, as increased sensitivity is of limited value if accompanied by numerous false-positive detections. In our more complex datasets, this trade-off was evident. PathoScope, Strainify and StrainR2 maintained high sensitivity but showed lower specificity and precision due to numerous low-abundance false-positive detections. Applying *post-hoc* detection thresholds mitigated this effect, improving tool specificity, precision and F1 score across both datasets. StrainR2 benefited particularly strongly, achieving the highest F1 score observed across our experiments (0.995). It also achieved the lowest mean absolute proportional error (0.06), consistent with its intended use-case for accurate strain abundance quantification[24]. However, the characteristics of the raw output remain an important practical consideration, as filtering numerous low-abundance false positives risks also removing genuine low-abundance strains. Indeed, at higher detection thresholds, sensitivity decreased for PathoScope and Strainify as some true-positive predictions were removed (Fig.6b). PanTax and StrainScan ran in super-low-depth mode offered a more conservative balance, maintaining high F1 scores without requiring *post-hoc* filtering. Tool selection should therefore consider not only overall performance, but also whether downstream filtering is appropriate for the intended application.

This distinction becomes important in real-world metagenomic profiling, where the strains present are unknown and may not be in the reference database. Under these conditions, assignment to a closely related reference, or even the correct broader phylogroup, would remain useful. Our experiments demonstrated that PathoScope and StrainGE both retained the underlying phylogenetic signal while frequently assigning strain identities to the closest available genomic neighbours to the ground truth by ANI. However, they achieved this through markedly different outputs. PathoScope more accurately reproduced the expected phylogroup-level abundances but produced more variable predictions requiring *post-hoc* filtering. StrainGE was less complete, failing to recover the low abundance phylogroup F, but consistently produced only one representative strain per detected phylogroup, providing a more conservative and readily interpretable output for downstream analysis. Nevertheless, our results support the use of either StrainGE or PathoScope for samples with unknown strain content, with selection depending on whether greater abundance accuracy or a more conservative, readily interpretable output is prioritised. These results suggest that paired-tool approaches may be advantageous when profiling “unknown” metagenomic samples. For example, PathoScope or StrainGE could first identify candidate phylogroups or closely related reference strains utilising a broad reference database. This could be followed by a tool with accurate strain-level abundance estimation (e.g., StrainR2) applied to the reduced search space. This approach could combine broad initial profiling with more targeted strain-level quantification, although this workflow was not evaluated here.

Precise strain-level resolution may not always be necessary for *E. coli* specifically, as broader phylogenetic classification can itself already be biologically informative in real-world applications. *E. coli* phylogroup composition already has clinical relevance, with phylogroup A more commonly associated with commensal lineages and B2 frequently associated with extraintestinal pathogenic *E. coli*[36–38]. Within clinical metagenomics applications[15], identifying the presence of a clinically relevant non-dominant phylogroup alongside other phylogroups could enhance useful clinical information compared to conventional routine-culture practices, even when the exact strains cannot be resolved with 100% certainty. Indeed, precise strain identity may only be clinically relevant where strains differ in consequential phenotypes, such as virulence or antibiotic resistance. Nevertheless, our results demonstrate that such phylogroup-level information can be retained by some tools despite absence of the true strain from the reference database.

For reference-based tools, strain-level profiling outputs are dependent on the contents of the reference genome database. This warrants consideration when interpreting previous applications of strain-level profiling to samples of unknown composition. For example, StrainGE has previously been applied to investigate strain dynamics before and after international travel[39], reporting 447 *E. coli* strain detections across 534 metagenomic samples (mean 0.5 strains pre-travel; 1.2 post-travel), and acquisition of phylogenetically distinct strains in 67% of travellers. The precise identities attributed to these detections are necessarily constrained by the 361-strain reference database used and cannot be assumed to represent exact matches to the biological strains present in each sample. Instead, they should be inferred to represent closely related reference strains. This does not challenge the evidence for strain turnover during international travel but instead highlights the distinction between detecting a phylogenetically distinct lineage and assigning its precise strain identity, with the latter inherently dependent on both tool resolution and reference database composition.

Interestingly, rapidly expanding reference databases may not improve strain-level profiling, as increasing numbers of highly similar genomes introduce redundancy and ambiguity in read assignment. Recent benchmarking found no correlation between reference genome database size and improved abundance accuracy or F1 score at strain-level, while targeted reference selection using hierarchical clustering improved profiling accuracy for highly similar strains[40]. Furthermore, our computational benchmarking highlights the practical constraints imposed by increasingly complex reference databases. Using 304 strains, StrainScan and PanTax already had peak memory requirements beyond those accessible without high-performance computing infrastructure, with PanTax being particularly resource intensive. Despite profiling with eight-fold more threads than the other tools, runtimes remained substantially longer, limiting practicality for large datasets. This presents an important trade-off, as PanTax, which supports the increasing availability of long-read metagenomic data[26], maintained high F1 scores across our datasets and its pangenome graph-based approach is designed to reduce reference redundancy while retaining strain-level diversity[41]. However, our results suggest the computational demands of this approach, including its underlying vg Giraffe short-read mapping framework[42], currently limit its accessibility and scalability.

Advantages of our study include the independent benchmarking of six competitive strain-level profiling tools across diverse metagenomic datasets, enabling identification of their advantages, disadvantages and suggested use cases (Table 2). We focused on *E. coli*, a common and genetically diverse member of the human gut microbiome and an important global One Health concern due to high rates of antibiotic resistance[43]. Although this focus may limit the transferability of our conclusions to other clinically relevant, strain-diverse gut colonisers with differing genomic characteristics, such as *Enterococcus faecium* and *Klebsiella pneumoniae*[44], we have provided the means to reproduce the benchmarking analysis and a framework within which to consider tool performance for other important bacterial species and/or microbiomes.

**Table 2.** Advantages, disadvantages and suggested use cases of the short-read metagenomic strain-level profiling tools evaluated in this study.

| Tool | Advantages | Disadvantages | Suggested use case(s) |
| --- | --- | --- | --- |
| StrainScan [22] | Comes with 8 pre-built reference genome databases, including *Escherichia coli*<br>Easy-to-use two module (strainscan_build & strainscan) command-line interface<br>Incorporated low depth modes (including super-low depth, benchmarked here) for rescued detection of low sequencing depth strains<br>High specificity and precision, regardless or detection threshold application | Reduced sensitivity at low sequencing depth<br>Low depth mode biases abundance estimate of the dominant strain leading to substantial overestimation<br>Out of phylogroup assignments if the true strain is removed from the reference database<br>High memory requirements and build time for larger reference databases<br>Can only profile one species at a time | Quick checks on a small number of metagenomic samples with a specific, user-desired reference database<br>For use on simulated metagenomes<br>For users with limited bioinformatics experience<br>For samples where a known, dominant strain is expected |
| Strainify [23] | Simple-to-use Nextflow workflow, including the ability to re-run the same database on new samples (--use_precomputed_variants parameter)<br>High sensitivity across datasets<br>Actively maintained in the last year (e.g., rewritten from Snakemake to Nextflow in v1.3.0)<br>Accessible runtime and memory requirements<br>Differentiates between strains at $\geq 99.9\%$ ANI across diverse metagenomic backgrounds | High false-positive detection rate (numerous low abundance false-positive detections)<br>Reports *E. coli* presence in negative controls<br>Detection thresholds ($\leq 0.005$) minimally improve specificity, precision and F1 score<br>Recent publication (October 2025), but yet to be peer-reviewed (at the time of writing) | Sensitivity applications (e.g., strain tracking) when the expected strain(s) in a set of given metagenome samples are known and a detection threshold can be applied<br>For users with limited bioinformatic experience |
| StrainR2 [24] | Easy-to-use two module (PreProcessR & StrainR) command-line interface<br>StrainR module provides a visualisation plot of FUKM abundances (unused in this study)<br>High F1 score tool (with detection threshold)<br>High accuracy tool for estimating strain-level relative abundance, in real-world samples<br>Differentiates between strains at $\geq 99.9\%$ ANI across diverse metagenomic backgrounds<br>Actively maintained in the last year<br>Accessible runtime and memory requirements | Application of detection thresholds to remove low abundance false positives when using complex reference databases may be an additional practical step/time limitation for some users<br>Requires the reference genomes of the strains present in the metagenome to provide accurate results | Highly accurate strain-level relative abundance predictions when expected strains in a set of given metagenome samples (simulated or real) are known (e.g., strain tracking)<br>Accurate strain-level abundance estimations using a reference database curated from a reduced phylogroup search space (paired tool usage approach)<br>For users with limited bioinformatics experience |
| **PathoScope 2.0** [25] | • Consistently high sensitivity and abundance estimation across datasets with minimal variation in predictions<br>• Historic tool (2014) which performs competitively relative to updated methodologies<br>• Well cited tool on PubMed (147 results)<br>• Accessible runtime and memory requirements | • Six modules but not all are inherently needed, leading to user-confusion e.g., requirement to self-generate Bowtie2 libraries (see Supplementary Methods)<br>• Ageing tool with no GitHub repository commits/updates since 2019<br>• Generation of large intermediate .sam files ($\geq 50$ Gb) during alignment steps with multiple references leading to storage concerns<br>• Reports *E. coli* presence in negative controls<br>• High false positive detection rate | • Sensitivity applications (e.g., strain tracking) when the expected strain(s) in a set of given metagenome samples are known and a detection threshold can be applied<br>• For accurate phylogroup-level detection on a novel dataset with unknown strain content (with a detection threshold e.g., 0.005 applied) |
| **PanTax** [26] | • Consistently high detection metrics and estimation accuracy performance across datasets<br>• Most modern tool benchmarked (Feb. 2026) and actively maintained<br>• Also works with long-read metagenomic data (not benchmarked in this study)<br>• Suitable for multi-species strain-level profiling (not benchmarked in this study) | • Inaccessible memory requirements outside of high-performance computation<br>• Long run times on complex samples<br>• Gurobi[48] optimizer necessary for performance but introduces user-restrictions and compatibility issues (see online Supplementary Material)<br>• Hard to install alongside Gurobi; bioinformatic experience required | • For accurate strain-level detection and abundance estimation in a high-performance computing environment<br>• Accurate strain-level abundance estimations using a reference database curated from a reduced phylogroup search space |
| **StrainGE** [27] | • Maintains within-phylogroup level assignments when true strains absent from the reference database and consistently reports “nearest neighbour” reference database strains<br>• Easy-to-follow documentation (see Supplementary Methods) with option to progress to strain-aware variant calling and nucleotide-level information using StrainGR (unused in this study)<br>• Well cited tool on PubMed (66 results), including previous *E. coli* strain-level diversity applications across a range of metagenomic studies<br>• Accepts a genus-wide reference database (multi-species strain profiling)<br>• Accessible runtime and memory requirements | • Relatively low strain-level abundance estimation accuracy across datasets<br>• Reduced sensitivity in complex datasets<br>• StrainGST and StrainGR are not implemented as a single, unified pipeline, requiring multiple sequential commands across modules, increasing workflow complexity<br>• Memory usage is variable dependent on paired metagenome file size | • For accurate phylogroup-level detection on a novel dataset with unknown strain content<br>• Followed with a more accurate abundance estimation tool (e.g., StrainR2) once a reduced phylogroup search space has been established (paired tool usage approach) |

Tool performance will inevitably evolve as existing methods are updated and new approaches merge. This is particularly relevant to long-read metagenomics, for which dedicated methods, including Strainberry[45] and MADRe[46], already exist. Continued benchmarking from the metagenomics community will be essential to monitor these developments. Finally, we note that our analysis focused exclusively on reference-based tools to ensure a consistent and comparable evaluation. The approaches assessed here are inherently limited in their ability to detect novel or previously uncharacterised *E. coli* strains lacking reference genomes, a limitation that *de novo* strain-level profiling methods can overcome[47].

## 9. Conclusions

This study represents the most recent benchmarking of short-read metagenomic strain-level profiling tools and the first to focus exclusively on *Escherichia coli*, a species exhibiting substantial strain-level diversity within the human gut. Across benchmarking datasets, tools produced markedly different representations of strain composition from the same metagenomic samples. We summarise the advantages and disadvantages of each tool and suggest the use of different methods depending on the intended application. Our findings will enable accurate characterisation of *E. coli* phylogroup- and strain-level diversity within short-read gut metagenomes in greater depth and at a larger-scale than achieved previously. This will advance our understanding of whether consistent genetic distinctions exist between commensal and pathogenic *E. coli* strains, with potential implications for the development of targeted therapeutics and vaccines.

## Supporting information

Supplementary Material version 2

## 10. Author statements

### 10.1 Author contributions

Matthew Galbraith: Data curation; Formal analysis; Investigation; Methodology; Software; Validation; Visualization, Writing – original draft.

David Williams: Supervision; Methodology; Writing – review & editing Liam P. Shaw: Supervision; Methodology; Writing – review & editing

Samuel Lipworth: Conceptualisation, Supervision; Methodology; Writing – review & editing

Nicole Stoesser: Conceptualisation, Funding acquisition, Methodology, Project administration, Resources, Supervision, Writing – review & editing

## 10.3 Funding information

This study/research is supported/funded by the National Institute for Health and Care Research (NIHR) Health Protection Research Unit: Healthcare Associated Infections and Antimicrobial Resistance (NIHR207397), a partnership between the UK Health Security Agency (UKHSA) and the University of Oxford. This work was also supported by the NIHR Oxford Biomedical Research Centre (BRC). Computation used the Oxford Biomedical Research Computing (BMRC) facility, a joint development between the Wellcome Centre for Human Genetics and the Big Data Institute supported by Health Data Research UK and the NIHR Oxford Biomedical Research Centre. The views expressed are those of the authors and not necessarily those of the NIHR, UKHSA or the Department of Health and Social Care.

Company Limited by Guarantee l Registered in England No. 1039582 l Registered Office Charles Darwin House, 12 Roger Street, WC1N 2JU, UK Registered as a Charity: 264017 (England & Wales); SC039250 (Scotland)

## 1.5 Repositories

N/A

