## Supplementary Material version 2 for "Benchmarking strain-level profiling of *Escherichia coli* in short-read gut metagenomes"

### Supplementary material for “Benchmarking strain-level profiling of *Escherichia coli* in short-read gut metagenomes” (Galbraith *et al*.,).

#### Supplementary Methods

##### S1. Example commands used per tool across benchmarking datasets

Example commands used to run each of the tools used in the benchmarking study can be found below. For full SLURM-scheduled job scripts used in the simulated gut metagenomes dataset and bash scripts used in the real-world spike-ins dataset (including computational benchmarking commands), see the Data Summary provided. For further information on individual tools, see their respective GitHub repositories (each linked below).

1.1). PanTax [1] v2.1.0 (https://github.com/LuoGroup2023/PanTax)

- Pre-step: Gurobi[2] v11.0.3 (academic license) was configured by setting environment variables (e.g. *GUROBI_ROOT*, *LD_LIBRARY_PATH*, and *PATH*) to the local installation directory, ensuring that the *gurobi_cl* executable was accessible during PanTax runs.
- Database construction:
  - pantax -f $genome_info --create
- Index database:
  - pantax -f $genome_info --db pantax_db --index
- Query database at species- and strain-level (short-reads, paired-end):
  - pantax -f $genome_info -s -p -r $R1 -r $R2 --db $pantax_db --species --strain
- When using PanTax v2.1.0, we also found the below helpful:
  - Note that the Gurobi academic license requires a one-time internet access for validation during the strain-level profiling step. This license is restricted to one device, making that step unsuitable for e.g., SLURM-scheduling on a high-performance computing cluster. See https://www.gurobi.com/academics for further information.
  - Based on the above, running PanTax during computational benchmarking within the real-world spike-ins dataset required a two-part script. Species-level profiling was performed via a SLURM-scheduled script employing the --next and --T parameters. Strain-level profiling was then performed on the dedicated compute node associated with our academic Gurobi license, again using the --next and --T parameters. This allowed the species-level profiling step, which includes the intensive vg Giraffe[3] pangenome read-mapping step that prevented PanTax reliably finishing on 4 threads, to be performed using a high number of threads (-t 32) without compromising strain-level profiling. For further information, see the Data Summary provided.

1.2). PathoScope [4] v2.0.7 (https://github.com/PathoScope/PathoScope)

- PathoMap module:
    - pathoscope MAP -1 $R1 -2 $R2 -targetRefFiles $REFS -outDir $mapOut -outAlign $.sam -expTag $dataset

    Where $REFS is a concatenated .fasta file of all reference genomes, for which a Bowtie2 index (including all .bt2 and .rev.*.bt2 files) was generated.
- PathoID module:
    - Pathoscope ID -alignFile $mapOut/*.sam -fileType $sam -outDir $idOut -expTag $dataset

    Note that PathoDB, PathoQC and PathoReport (optional modules) were unused. PathoLib (core module) was also unused, as we manually curated our reference genome databases for all datasets and concatenated/built a Bowtie2 index independently (see above).

1.3). StrainGE [5] v1.3.9 (https://github.com/broadinstitute/StrainGE & https://strainge.readthedocs.io/en/latest/)

- Kmerize reference genomes:
    - for f in $strainge_db/*.fa.gz; do; straingst kmerize -k31 -o ${f%.fa.gz}.hdf5 $f; done
- Create pangenome k-mer database:
    - straingst createdb -o $pangenome_db $strainge_db/*.hdf5
- Kmerize samples
    - straingst kmerize -k 31 -o $sample.hdf5 $R1 $R2
- Run StrainGST
    - straingst run --separate-output -o $sample $pangenome_db $sample.hdf5

  Note that the StrainGR module was not used for estimated relative abundances as the developers advise using abundances calculated by StrainGST.

1.4). Strainify [6] v1.3.1 (https://github.com/treangenlab/Strainify)

- Snakemake Nextflow workflow launched via:
    - strainify --genome_folder $genome_dir --fastq_folder $sample_dir --outdir $outroot
- If multiple samples were run on the same database, the use precomputed variants option was used:
    - strainify --genome_folder $genome_dir --fastq_folder $sample_dir --use_precomputed_variants --precomputed_dir $precomp_dir --outdir $outroot

1.5). StrainR2 [7] v2.3.0 (https://github.com/BisanzLab/StrainR2)

- Genome preprocessing and database creation using PreProcessR:
    - PreProcessR -I $reference_genome_database -o $StrainR2_db
- Profiling and abundance estimation using StrainR:
    - StrainR -1 $R1 -2 $R2 -r $StrainR2_db -o $sample_results

1.6). StrainScan [8] v1.0.14 (https://github.com/liaoherui/StrainScan)

- Build custom database:
  - strainscan_build -i $reference_genome_database -o $strainscan_db
- Build custom database using the memory efficient mode:
  - strainscan_build -i $reference_genome_database -o $strainscan_db -e 1
- Identify bacterial strains in short-reads:
  - strainscan -i $R1 -j $R2 -d $strainscan_db -o $sample_results
- Identify bacterial strains in short-reads using StrainScan super low depth mode, with probabilistic strain detection enabled:
  - strainscan -i $R1 -j $R2 -d $strainscan_db -l 2 -b 1 -o $sample_results

#### S2. Construction of benchmarking datasets

Detailed information on the construction and processing of each of the four benchmarking datasets can be found below.

*(i) ZymoBIOMICS® D6331 gut microbiome standard*

Genome assemblies of the five D6331 *E. coli* strains were obtained from Zymo Research (https://s3.amazonaws.com/zymo-files/BioPool/D6331.refseq.zip). Phylogroup assignment and sequence typing were performed using EzClermont[9] v0.6.3 and mlst v2.23.0[10] with the Achtman E. coli PubMLST scheme[11], respectively. A previously generated simulated paired-end 150 bp short-read dataset (ShortPE_mock; SRR25211204) was downloaded from the NCBI Sequence Read Archive; this was generated from the original long-read (PacBio) sequencing dataset of the D6331 standard (Zymo_D6331; SRS7763867). Although residual sequencing errors may therefore be inherited by the simulated short reads, the source HiFi reads were generated using circular consensus sequencing (CCS) with a minimum predicted accuracy of 99%[12] and are unexpected to affect strain-level profiling here.

*(ii) SRR13355226 mock community*

SRR13355226 had human reads removed using Deacon[13] v0.13.2, utilising the 3 GB human pangenome index and default parameters. This generated paired-end FASTQ files depleted of human reads, which we have made publicly available (SRR13355226_depleted; see Data summary).

*(iii) Simulated gut metagenomes*

*E. coli* relative abundances in GutFeelingKB[14] were aggregated into a single species entry by summing the abundances of the O104:H4 (0.0004) and O83:H1 (0.0006) strains with the dominant *E. coli* species entry (0.0187). The remaining 107 species were cross-referenced against the NCBI Genome database (https://www.ncbi.nlm.nih.gov/datasets/genome/) to identify available whole-genome assemblies, prioritising RefSeq and single-contig assemblies. We excluded nine species due to the absence of complete genome assemblies (*Ruminococcus champanellensis*, *Coprococcus catus*, *Gordonibacter pamelaeae*, butyrate-producing bacterium SM4/1, butyrate-producing bacterium SS3/4, *Streptococcus* sp. I-P16, *Fermentimonas caenicola*, *Ruminococcus bromii*, and *Faecalitalea cylindroides*), leaving 98 assemblies. Relative abundances were normalised, with abundances distributed across contigs in proportion to contig length for multi-contig assemblies (see Data Summary). The final *E. coli* relative abundance (0.0196) was split between two strain-level NCBI reference genome assemblies (K-12 substr. MG1655, GCF_000005845.2; O157:H7 str. Sakai, GCF_000008865.2) across a range of abundance ratios (0.50:0.50, 0.75:0.25, 0.90:0.10, 0.95:0.05, 0.99:0.01). For the negative control (0.00:0.00), *E. coli* was removed and relative abundances renormalised across the remaining 97 species. These files, together with the identified genome assemblies, were used as input for metagenome simulation with

InSilicoSeq[15], using the HiSeq Illumina 150bp paired-end error model across sequencing efforts of 20, 50, and 100 million paired-end reads, and in fixed-seed triplicates (--seed 1/2/3). Simulation scripts and all abundance files describing per-contig, per-species relative abundances are provided (see Data Summary).

Phylogroups within the 24-strain *E. coli* reference database were confirmed using EzClermont[9] v0.6.3, except for phylogroup C which was based on previously defined genome-wide Mash clustering[16]. Pairwise ANI were calculated using FastANI[17] v1.34, comparing K-12 substr. MG1655 and O157:H7 str. Sakai against all other reference genomes (Table S6).

*(iv) Real-world spike-ins*

42 publicly available short-read gut metagenomes that had previously been identified to be absent for *E. coli* reads (relative abundance detection limit of 0.003–0.01%[18]) were selected. Identified metagenomic read files were downloaded and quality controlled using the metagen-fetch script published by the same authors (https://github.com/alexmsalmeida/metagen-fetch). Downloaded metagenomic runs from the same sample were concatenated to produce paired-end read files per sample. Kraken2 [19] v2.17.1 was run using the standard database (https://genome-idx.s3.amazonaws.com/kraken/k2_standard_20260626.tar.gz) on all metagenome samples to identify missed low abundance *E. coli* reads. Reads assigned to *E. coli* were subsequently removed using the extract_reads.py script from KrakenTools[20] v1.2.1 (https://github.com/jenniferlu717/KrakenTools), utilising the --tax_id 562 and --include_children parameters. Kraken2 was rerun to ensure the metagenomes were depleted of *E. coli* reads.

To generate the 304-strain *E. coli* reference database, we identified a set of 10,667 *E. coli* genomes[16]. Following *Shigella* removal, 9,384 *E. coli* genomes remained, of which 9,382 were downloadable from NCBI. Assembly statistics were obtained using NCBI Datasets v18.26.0[21] and merged with the original dataset. This was filtered for complete genomes, to avoid potential contamination from draft genomes during strain-profiling[20]. 602 complete genomes were retained and assessed across CheckM completeness and contamination thresholds to balance genome quality, dataset size, and phylogroup representation (see Data Summary). Final thresholds of ≥97% completeness and ≤3% contamination were applied, leaving 526 complete genomes for download. These were dereplicated using dRep[22] v3.6.1, using primary (-pa) and secondary ANI thresholds (-sa) of 0.90 and 0.999 respectively, incorporating the --S_algorithm fastANI and --ignoreGenomeQuality paramaters. This produced a final *E. coli* reference database comprising 304 strains for input to all tools. This, alongside an R script to reproducibly generate this database, are available (see Data summary).

##### S3. Performance metrics equations

The full equations used to calculate the estimation and detection metrics are provided below (Eqs. S2.1–S2.10). All variables and parameters are defined following the equations. Note that absolute proportional error is undefined when $p_i = 0$. Absolute proportional error can exceed 1 when predicted abundances deviate substantially from low true abundances, whereas missed strain detections (predicted abundance = 0) are assigned an absolute proportional error of 1.

Let:

- $p_i$= true relative abundance of strain $i$
- $\hat{p}_i$= predicted relative abundance of strain $i$
- $N$= total number of strains

$$\text{Error}_i = \hat{p}_i - p_i \quad \text{(Eq. S2.1)}$$

$$\text{Absolute Error}_i = |\hat{p}_i - p_i| \quad \text{(Eq. S2.2)}$$

$$\text{Absolute Proportional Error}_i = \frac{|\hat{p}_i - p_i|}{p_i} \quad \text{(Eq. S2.3)}$$

$$\text{Mean Error} = \frac{1}{N}\sum_{i=1}^{N}(\hat{p}_i - p_i) \quad \text{(Eq. S2.4)}$$

$$\text{Mean Absolute Error} = \frac{1}{N}\sum_{i=1}^{N}|\hat{p}_i - p_i| \quad \text{(Eq. S2.5)}$$

$$\text{Mean Absolute Proportional Error} = \frac{1}{N}\sum_{i=1}^{N}\frac{|\hat{p}_i - p_i|}{p_i} \quad \text{(Eq. S2.6)}$$

$$\text{Sensitivity} = \frac{True\ Positives}{True\ Positives + False\ Negatives} \quad \text{(Eq. S2.7)}$$

$$\text{Specificity} = \frac{True\ Negatives}{True\ Negatives + False\ Positives} \quad \text{(Eq. S2.8)}$$

$$\text{Precision} = \frac{True\ Positives}{True\ Positives + False\ Positives} \quad \text{(Eq. S2.9)}$$

$$\text{F1 score} = 2 * \frac{Precision * Sensitivity}{Precision + Sensitivity} \quad \text{(Eq. S2.10)}$$

##### S4. R packages

The following R packages were used with R[23] v4.5.1 as part of the statistical analysis and visualisation of this work: readxl[24] v1.4.5, ggplot2[25] v4.0.0, cowplot[26] v1.2.0, dplyr[27] v1.1.4, stringr[28] v1.6.0, tidyr[29] v1.3.1, forcats[30] v1.0.1, scales[31] v1.4.0, binom[32] v1.1.1.1 and purrr[33] v1.1.0, ggbreak[34], RColorBrewer[35] v1.3-3, colorspace[36] v2.1.2, ggtext[37] v0.2.0 and ggplotify[38] v0.1.3. Full R scripts for reproducible analysis have been made available (see Data Summary).

### Supplementary Tables

**Table S1. Identified short-read metagenomic strain-level analysis tools excluded from this study.** *as of 05/10/2026.

| Tool reference | Primary approach | Installation | Version trialled (last commit)* | Reason for exclusion |
|---|---|---|---|---|
| **StrainEst [39]** | Single nucleotide variants profiles of the user-provided reference genomes to identify strains | Docker; GitHub repository (clone source); Bioconda | v1.2.4 (Jan 17, 2020) | Limited maintainability and database construction challenges; performance reported to be consistently lower than more recent methods[8] |
| **StrainFLAIR [40]** | Strain identification and quantification using variation graphs representative of gene sequences | GitHub repository (clone source) | v.0.0.1 (Jan 29, 2023) | Tool no longer supported and software no longer maintained since March 31st, 2022. |
| **MixtureS [41]** | Reference-guided *de novo* strain reconstruction using identified polymorphic sites from mapped reads | UCF project webpage | v1.0 (publication date: Aug 17, 2020) | Primary output is an inferred haplotype. Lack of public (e.g., GitHub) repository. |
| **kSanity [42]** | Reference-based k-mer indexing approach, leveraging strain-unique and shared k-mers to detect and quantify strains based on their relative representation in metagenomic sequencing data. | GitHub repository (clone source) | No formal release (GitHub main, latest commit: Aug 20, 2025) | Primary emphasis is on identifying one focal strain from a conspecific reference database and longitudinal strain tracking. Lack of reproducible version information available for install. |
| **MetaPhlAn, StrainPhlAn [43, 44]** | SNV-based reconstruction of consensus strain sequences from species-specific marker genes. | Bioconda; GitHub repository (clone source) | v4.2.4 (Dec 13, 2025) | Designed for dominant strain reconstruction and cross-sample phylogenetic comparison as part of the Biobakery 3 package, rather than within-sample multi-strain resolution. |
| **PanPhlAn [45]** | Pangenome-based profiling that infers strain gene presence /absence from metagenomic read mapping. | Bioconda; GitHub repository (clone source); pip install | v3.1 (Oct 3, 2023) | Infers strain gene-content profiles without directly estimating relative strain abundances. |

| | | | | |
|---|---|---|---|---|
| **Sigma [46]** | Reference genome alignment with maximum-likelihood estimation for strain identification and abundance estimation. | GitHub repository (manual source download and compilation) | v1.1.0 (Oct 17, 2023) | Not included due to practical challenges in installation and workflow configuration. |
| **ConStrains [47]** | Reconstructs strain diversity and SNPs by deconvoluting SNP patterns across different conserved genes and samples | Bitbucket repository (install from source) | No formal release (Bitbucket master branch) | Requires high sequencing coverage and complex SNP-based strain inference, which limited practical inclusion in this benchmarking framework. |
| **StrainPRO [48]** | Identifies representative sequence segments from user-provided reference genomes to characterise strain-level profiles and relative abundances | GitHub repository (clone source) | v0.9.8 (July 9, 2020) | Not peer reviewed since BioRxiv deposition in 2019. |
| **Floria [49]** | Minimum error correction read clustering and strain-preserving network flow model on short and long-read metagenomic sequencing data | GitHub repository (clone source); Bioconda | v0.0.2 (Apr 11, 2025) | Outputs number of strains per sample but lacks comparative relative abundance predictions. Only allows one reference genome as alignment input. |
| **STRONG [50]** | Reconstructs strain genomes *de novo* by leveraging co-assembly graphs across samples and using Bayesian inference to resolve strain-specific paths and abundances | GitHub repository (clone source) + run install script | v0.0.1-beta (Apr 13, 2023) | Focuses on *de novo* strain assembly using reference-free based methods. |
| **QMatey [51]** | MegaBLAST alignment / diagnostic-read exact matching | GitHub; Bash/R pipeline with external dependencies | v0.5.2 (Oct 26, 2025) | Unable to reproduce a successful strain-profiling analysis using the publicly available software and documentation despite repeated attempts. Limited evidence of independent uptake in the published literature since its release in 2023. |

**Table S2. ZymoBIOMICS® D6331 gut microbiome standard theoretical genomic DNA compositions (%), with *Escherichia coli* phylogroups and sequence types.**

| Species (strain) | Genomic DNA (%) | *Escherichia coli* scaled relative abundance | Phylogroup (*E. coli* only) | Sequence type (ST; *E. coli* only) | Allelic ST profile (*E. coli* only; Achtman scheme) |
|---|---|---|---|---|---|
| *Faecalibacterium prausnitzii* | 14 | NA | NA | NA | NA |
| *Veillonella rogosae* | 14 | NA | NA | NA | NA |
| *Roseburia hominis* | 14 | NA | NA | NA | NA |
| *Bacteroides fragilis* | 14 | NA | NA | NA | NA |
| *Prevotella copri* | 6 | NA | NA | NA | NA |
| *Bifidobacterium adolescentis* | 6 | NA | NA | NA | NA |
| *Fusobacterium nucleatum* | 6 | NA | NA | NA | NA |
| *Lactobacillus fermentum* | 6 | NA | NA | NA | NA |
| *Clostridioides difficile* | 1.5 | NA | NA | NA | NA |
| *Akkermansia muciniphila* | 1.5 | NA | NA | NA | NA |
| *Methanobrevibacter smithii* | 0.1 | NA | NA | NA | NA |
| *Salmonella enterica* | 0.01 | NA | NA | NA | NA |
| *Enterococcus faecalis* | 0.001 | NA | NA | NA | NA |
| *Clostridium perfringens* | 0.0001 | NA | NA | NA | NA |
| *Escherichia coli (JM109)* | 2.8 | 0.2 | A | 10 | adk(10), fumC(11), gyrB(4), icd(8), mdh(8), purA(8), recA(2) (ST 10) |
| *Escherichia coli (B-3008)* | 2.8 | 0.2 | A | 3021 | adk(8), fumC(7), gyrB(4), icd(8), mdh(8), purA(18), recA (233) (ST 3021) |
| *Escherichia coli (B-2207)* | 2.8 | 0.2 | EC control fail | 7610 | adk(10), fumC(11), gyrB(4), icd(852), mdh(8), purA(8), recA(2) (ST 7610) |
| *Escherichia coli (B-766)* | 2.8 | 0.2 | B1 | novel | adk(6), fumC(19), gyrB(14), icd(16), mdh(11), purA(12), recA(2) (ST 1079) |
| *Escherichia coli (B-1109)* | 2.8 | 0.2 | A | 10 | adk(10), fumC(11), gyrB(4), icd(8), mdh(8), purA(8), recA(2) (ST10) |
| *Candida albicans* | 1.5 | NA | NA | NA | NA |
| *Saccharomyces cerevisiae* | 1.4 | NA | NA | NA | NA |

**Table S3. Composition of *Escherichia coli* strains within the SRR13355226 mock community (99% human, 1% *E. coli* DNA; 6,480,879,230 bp; 21,459,865 spots).** Estimated sequencing depth (x) for each strain was calculated as: ((total metagenomic bases x sensitivity x ground-truth strain proportion)/reference genome size).

| ***Escherichia coli* strain** | **Genome accession** | ***E. coli* scaled relative abundance** | **Genome size (Mb)** | **Approximate bases from strain (bp)** | **Spots** | **Approximate reads** | **Estimated sequencing depth (x)** |
|---|---|---|---|---|---|---|---|
| H10407 | GCF_000210475.1 | 0.8 | 5.3 | 51,847,033.80 | 171,673 | 343,346 | 9.79 |
| UTI89 | GCF_015644765.1 | 0.15 | 5.1 | 9,721,318.80 | 32,190 | 64,380 | 1.91 |
| O157:H7 str. Sakai | GCF_000008865.2 | 0.049 | 5.6 | 3,175,631.80 | 10,515 | 21,030 | 0.57 |
| O139:H28 str. E24377A | GCF_000017745.1 | 0.001 | 5.2 | 64,808.80 | 215 | 430 | 0.012 |

**Table S4. Final InSilicoSeq setup for metagenome simulation.** *K12-MG1655 : O157:H7 str. Sakai ratio reflects the imposed relative abundance ratio between the two *Escherichia coli* strains. ‡Relative-abundance values for K12-MG1655 and O157:H7 str. Sakai refer to their proportions within the entire simulated metagenome.

| Total reads (--n_reads) | True read pairs (sequencing effort) | K12-*MG1655*: *O157:H7* str. Sakai E. coli ratio* | K12-MG1655 relative abundance‡ | K12 -MG1655 simulated sequencing depth (×) | O157:H7 str. Sakai relative abundance‡ | O157:H7 str. Sakai simulated sequencing depth (×) |
|---|---|---|---|---|---|---|
| 40000000 | 20000000 | 50:50 | 0.0098 | 12.66790358 | 0.0098 | 10.51012538 |
| 40000000 | 20000000 | 75:25 | 0.0147 | 19.00185537 | 0.0049 | 5.25506269 |
| 40000000 | 20000000 | 90:10 | 0.01764 | 22.80222645 | 0.00196 | 2.102025076 |
| 40000000 | 20000000 | 95:5 | 0.01862 | 24.06901681 | 0.00098 | 1.051012538 |
| 40000000 | 20000000 | 99:1 | 0.019404 | 25.08244909 | 0.000196 | 0.210202508 |
| 40000000 | 20000000 | 0:0 | 0 | 0 | 0 | 0 |
| 100000000 | 50000000 | 50:50 | 0.0098 | 31.66975896 | 0.0098 | 26.27531345 |
| 100000000 | 50000000 | 75:25 | 0.0147 | 47.50463843 | 0.0049 | 13.13765672 |
| 100000000 | 50000000 | 90:10 | 0.01764 | 57.00556612 | 0.00196 | 5.25506269 |
| 100000000 | 50000000 | 95:5 | 0.01862 | 60.17254202 | 0.00098 | 2.627531345 |
| 100000000 | 50000000 | 99:1 | 0.019404 | 62.70612273 | 0.000196 | 0.525506269 |
| 100000000 | 50000000 | 0:0 | 0 | 0 | 0 | 0 |
| 200000000 | 100000000 | 50:50 | 0.0098 | 63.33951791 | 0.0098 | 52.5506269 |
| 200000000 | 100000000 | 75:25 | 0.0147 | 95.00927687 | 0.0049 | 26.27531345 |
| 200000000 | 100000000 | 90:10 | 0.01764 | 114.0111322 | 0.00196 | 10.51012538 |
| 200000000 | 100000000 | 95:5 | 0.01862 | 120.345084 | 0.00098 | 5.25506269 |
| 200000000 | 100000000 | 99:1 | 0.019404 | 125.4122455 | 0.000196 | 1.051012538 |
| 200000000 | 100000000 | 0:0 | 0 | 0 | 0 | 0 |

**Table S5. 24-strain *Escherichia coli* reference genome database used as input for the simulated gut metagenomes dataset.** Three strains from each of the eight major phylogroups were selected for reference database inclusion. Genomes were downloaded from NCBI (https://www.ncbi.nlm.nih.gov/datasets/genome/) using ncbi-genome-download v0.3.3[52]. K12-MG1655 and O157:H7 str. Sakai were excluded from the reference database for the simulated metagenomes reduced reference database experiments.

| Strain | Genome accession | Phylogroup |
|---|---|---|
| K-12 MG1655 | GCF_000005845.2_ASM584v2_genomic | A |
| HS | GCF_000017765.1_ASM1776v1_genomic | A |
| REL606 | GCF_000017985.1_ASM1798v1_genomic | A |
| SE11 | GCF_000010385.1_ASM1038v1_genomic | B1 |
| O139:H28 str. E24377A | GCF_000017745.1_ASM1774v1_genomic | B1 |
| IAI1 | GCF_000026265.1_ASM2626v1_genomic | B1 |
| UTI89 | GCF_000013265.1_ASM1326v1_genomic | B2 |
| 536 | GCF_000013305.1_ASM1330v1_genomic | B2 |
| IHE3034 | GCF_000025745.1_ASM2574v1_genomic | B2 |
| O26:H11 str.11368 | GCF_000091005.1_ASM9100v1_genomic | C |
| 97-3250 | GCF_003018455.1_ASM301845v1_genomic | C |
| FORC_028 | GCF_001721125.1_ASM172112v1_genomic | C |
| UMN026 | GCF_000026325.1_ASM2632v2_genomic | D |
| O42 | GCF_000027125.1_ASM2712v1_genomic | D |
| PCN033 | GCF_000219515.2_ASM21951v3_genomic | D |
| O157:H7 str. Sakai | GCF_000008865.2_ASM886v2_genomic | E |
| O55:H7 str. CB9615 | GCF_000025165.1_ASM2516v1_genomic | E |
| O157:H7 str. EDL933 | GCF_000732965.1_ASM73296v1_genomic | E |
| SMS-3-5 | GCF_000019645.1_ASM1964v1_genomic | F |
| IAI39 | GCF_000026345.1_ASM2634v1_genomic | F |
| 2016C-3878 | GCF_003204955.1_ASM320495v1_genomic | F |
| 2009C-3133 | GCF_001420955.1_ASM142095v1_genomic | G |
| 2014C-3338 | GCF_003018095.1_ASM301809v1_genomic | G |
| 2012C-4502 | GCF_003018255.1_ASM301825v1_genomic | G |

**Table S6. FastANI pairwise average nucleotide identity of K12-MG1655 and O157:H7 str. Sakai to all reference genomes used in the InSilicoSeq-simulated metagenomes dataset.** All reference genomes can be found in Table S5, above. Count of bidirectional fragment mappings and total query fragments not shown.

| Query genome accession | Query genome strain | Query genome phylogroup | Reference genome accession | Reference genome strain | Reference genome phylogroup | Average nucleotide identity (%) |
|---|---|---|---|---|---|---|
| GCF_000005845.2_ASM584v2_genomic.fna | K12 MG1655 | A | GCF_000017985.1_ASM1798v1_genomic.fna | REL606 | A | 98.951 |
| GCF_000005845.2_ASM584v2_genomic.fna | K12 MG1655 | A | GCF_000017765.1_ASM1776v1_genomic.fna | HS | A | 98.743 |
| GCF_000005845.2_ASM584v2_genomic.fna | K12 MG1655 | A | GCF_000026265.1_ASM2626v1_genomic.fna | IAI1 | B1 | 98.476 |
| GCF_000005845.2_ASM584v2_genomic.fna | K12 MG1655 | A | GCF_000010385.1_ASM1038v1_genomic.fna | SE11 | B1 | 98.463 |
| GCF_000005845.2_ASM584v2_genomic.fna | K12 MG1655 | A | GCF_000017745.1_ASM1774v1_genomic.fna | O139:H28 str. E24377A | B1 | 98.395 |
| GCF_000005845.2_ASM584v2_genomic.fna | K12 MG1655 | A | GCF_003018455.1_ASM301845v1_genomic.fna | 97-3250 | C | 98.371 |
| GCF_000005845.2_ASM584v2_genomic.fna | K12 MG1655 | A | GCF_000091005.1_ASM9100v1_genomic.fna | O26:H11 str.11368 | C | 98.360 |
| GCF_000005845.2_ASM584v2_genomic.fna | K12 MG1655 | A | GCF_001721125.1_ASM172112v1_genomic.fna | FORC_028 | C | 98.331 |
| GCF_000005845.2_ASM584v2_genomic.fna | K12 MG1655 | A | GCF_000025165.1_ASM2516v1_genomic.fna | O55:H7 str. CB9615 | E | 97.865 |
| GCF_000005845.2_ASM584v2_genomic.fna | K12 MG1655 | A | GCF_000008865.2_ASM886v2_genomic.fna | O157:H7str. Sakai | E | 97.856 |
| GCF_000005845.2_ASM584v2_genomic.fna | K12 MG1655 | A | GCF_000732965.1_ASM73296v1_genomic.fna | O157:H7 str. EDL933 | E | 97.842 |
| GCF_000005845.2_ASM584v2_genomic.fna | K12 MG1655 | A | GCF_000027125.1_ASM2712v1_genomic.fna | O42 | D | 97.353 |
| GCF_000005845.2_ASM584v2_genomic.fna | K12 MG1655 | A | GCF_000026325.1_ASM2632v2_genomic.fna | UMN026 | D | 97.344 |
| GCF_000005845.2_ASM584v2_genomic.fna | K12 MG1655 | A | GCF_000219515.2_ASM21951v3_genomic.fna | PCN033 | D | 97.239 |
| GCF_000005845.2_ASM584v2_genomic.fna | K12 MG1655 | A | GCF_001420955.1_ASM142095v1_genomic.fna | 2009C-3133 | G | 97.207 |

| | | | | | | |
|---|---|---|---|---|---|---|
| GCF_000005845.2_ASM584v2_genomic.fna | K12 MG1655 | A | GCF_003204955.1_ASM320495v1_genomic.fna | 2016C-3878 | F | 97.190 |
| GCF_000005845.2_ASM584v2_genomic.fna | K12 MG1655 | A | GCF_003018095.1_ASM301809v1_genomic.fna | 2014C-3338 | G | 97.130 |
| GCF_000005845.2_ASM584v2_genomic.fna | K12 MG1655 | A | GCF_000019645.1_ASM1964v1_genomic.fna | SMS-3-5 | F | 97.081 |
| GCF_000005845.2_ASM584v2_genomic.fna | K12 MG1655 | A | GCF_000026345.1_ASM2634v1_genomic.fna | IAI39 | F | 97.072 |
| GCF_000005845.2_ASM584v2_genomic.fna | K12 MG1655 | A | GCF_003018255.1_ASM301825v1_genomic.fna | 2012C-4502 | G | 97.061 |
| GCF_000005845.2_ASM584v2_genomic.fna | K12 MG1655 | A | GCF_000013305.1_ASM1330v1_genomic.fna | 536 | B2 | 96.754 |
| GCF_000005845.2_ASM584v2_genomic.fna | K12 MG1655 | A | GCF_000025745.1_ASM2574v1_genomic.fna | IHE3034 | B2 | 96.711 |
| GCF_000005845.2_ASM584v2_genomic.fna | K12 MG1655 | A | GCF_000013265.1_ASM1326v1_genomic.fna | UTI89 | B2 | 96.687 |
| GCF_000008865.2_ASM886v2_genomic.fna | O157:H7 str. Sakai | E | GCF_000732965.1_ASM73296v1_genomic.fna | O157:H7 str. EDL933 | E | 99.969 |
| GCF_000008865.2_ASM886v2_genomic.fna | O157:H7 str. Sakai | E | GCF_000025165.1_ASM2516v1_genomic.fna | O55:H7 str. CB9615 | E | 99.471 |
| GCF_000008865.2_ASM886v2_genomic.fna | O157:H7 str. Sakai | E | GCF_000005845.2_ASM584v2_genomic.fna | K-12 MG1655 | A | 97.762 |
| GCF_000008865.2_ASM886v2_genomic.fna | O157:H7 str. Sakai | E | GCF_000026265.1_ASM2626v1_genomic.fna | IAI1 | B1 | 97.729 |
| GCF_000008865.2_ASM886v2_genomic.fna | O157:H7 str. Sakai | E | GCF_000017765.1_ASM1776v1_genomic.fna | HS | A | 97.681 |
| GCF_000008865.2_ASM886v2_genomic.fna | O157:H7 str. Sakai | E | GCF_000017985.1_ASM1798v1_genomic.fna | REL606 | A | 97.647 |
| GCF_000008865.2_ASM886v2_genomic.fna | O157:H7 str. Sakai | E | GCF_001721125.1_ASM172112v1_genomic.fna | FORC_028 | C | 97.623 |
| GCF_000008865.2_ASM886v2_genomic.fna | O157:H7 str. Sakai | E | GCF_003018455.1_ASM301845v1_genomic.fna | 97-3250 | C | 97.600 |
| GCF_000008865.2_ASM886v2_genomic.fna | O157:H7 str. Sakai | E | GCF_000091005.1_ASM9100v1_genomic.fna | O26:H11 str.11368 | C | 97.586 |
| GCF_000008865.2_ASM886v2_genomic.fna | O157:H7 str. Sakai | E | GCF_000017745.1_ASM1774v1_genomic.fna | O139:H28 str. E24377A | B1 | 97.516 |
| GCF_000008865.2_ASM886v2_genomic.fna | O157:H7 str. Sakai | E | GCF_000010385.1_ASM1038v1_genomic.fna | SE11 | B1 | 97.512 |

| | | | | | | |
|---|---|---|---|---|---|---|
| GCF_000008865.2_ASM886v2_genomic.fna | O157:H7 str. Sakai | E | GCF_000219515.2_ASM21951v3_genomic.fna | PCN033 | D | 97.107 |
| GCF_000008865.2_ASM886v2_genomic.fna | O157:H7 str. Sakai | E | GCF_000026325.1_ASM2632v2_genomic.fna | UMN026 | D | 96.929 |
| GCF_000008865.2_ASM886v2_genomic.fna | O157:H7 str. Sakai | E | GCF_003204955.1_ASM320495v1_genomic.fna | 2016C-3878 | F | 96.872 |
| GCF_000008865.2_ASM886v2_genomic.fna | O157:H7 str. Sakai | E | GCF_000026345.1_ASM2634v1_genomic.fna | IAI39 | F | 96.833 |
| GCF_000008865.2_ASM886v2_genomic.fna | O157:H7 str. Sakai | E | GCF_000019645.1_ASM1964v1_genomic.fna | SMS-3-5 | F | 96.821 |
| GCF_000008865.2_ASM886v2_genomic.fna | O157:H7 str. Sakai | E | GCF_003018095.1_ASM301809v1_genomic.fna | 2014C-3338 | G | 96.770 |
| GCF_000008865.2_ASM886v2_genomic.fna | O157:H7 str. Sakai | E | GCF_000027125.1_ASM2712v1_genomic.fna | O42 | D | 96.758 |
| GCF_000008865.2_ASM886v2_genomic.fna | O157:H7 str. Sakai | E | GCF_003018255.1_ASM301825v1_genomic.fna | 2012C-4502 | G | 96.742 |
| GCF_000008865.2_ASM886v2_genomic.fna | O157:H7 str. Sakai | E | GCF_001420955.1_ASM142095v1_genomic.fna | 2009C-3133 | G | 96.722 |
| GCF_000008865.2_ASM886v2_genomic.fna | O157:H7 str. Sakai | E | GCF_000013305.1_ASM1330v1_genomic.fna | 536 | B2 | 96.515 |
| GCF_000008865.2_ASM886v2_genomic.fna | O157:H7 str. Sakai | E | GCF_000025745.1_ASM2574v1_genomic.fna | IHE3034 | B2 | 96.458 |
| GCF_000008865.2_ASM886v2_genomic.fna | O157:H7 str. Sakai | E | GCF_000013265.1_ASM1326v1_genomic.fna | UTI89 | B2 | 96.402 |

**Table S7. Publicly available metagenome samples used in the real-world metagenomes multi-strain spike-ins dataset**. 42 sample-level metagenomes were selected based on previously curated data on publicly available short-read gut metagenomes [18]. Presented are run, sample and study-level metadata to allow reproducible download and quality control via the open-source metagen-fetch script produced by the same authors (https://github.com/alexmsalmeida/metagen-fetch). Runs were concatenated to the sample-level following completion of the script. Sample-level *E. coli* read counts were obtained from the estimated UHGG abundance read counts from the same study (https://figshare.com/articles/dataset/UHGG_abundance_read_counts/27044341/1?file=49238707 ). Instrument platform, instrument model and library layout were retrieved for each run accession from the European Nucleotide Archive (ENA) Portal API and incorporated into the master metadata table.

| Run | Sample | Study | Country | Continent | Sample-level *E. coli* read counts | Instrument platform | Instrument model | Library layout |
|---|---|---|---|---|---|---|---|---|
| **ERR479041** | ERS436645 | ERP005534 | France | Europe | 0 | ILLUMINA | Illumina HiSeq 2000 | PAIRED |
| **ERR479042** | ERS436645 | ERP005534 | France | Europe | 0 | ILLUMINA | Illumina HiSeq 2000 | PAIRED |
| **ERR479049** | ERS436647 | ERP005534 | France | Europe | 0 | ILLUMINA | Illumina HiSeq 2000 | PAIRED |
| **ERR479050** | ERS436647 | ERP005534 | France | Europe | 0 | ILLUMINA | Illumina HiSeq 2000 | PAIRED |
| **ERR479051** | ERS436647 | ERP005534 | France | Europe | 0 | ILLUMINA | Illumina HiSeq 2000 | PAIRED |
| **ERR479052** | ERS436647 | ERP005534 | France | Europe | 0 | ILLUMINA | Illumina HiSeq 2000 | PAIRED |
| **ERR479393** | ERS436655 | ERP005534 | France | Europe | 0 | ILLUMINA | Illumina HiSeq 2000 | PAIRED |
| **ERR479394** | ERS436655 | ERP005534 | France | Europe | 0 | ILLUMINA | Illumina HiSeq 2000 | PAIRED |
| **ERR479167** | ERS436658 | ERP005534 | France | Europe | 0 | ILLUMINA | Illumina HiSeq 2000 | PAIRED |
| **ERR479168** | ERS436658 | ERP005534 | France | Europe | 0 | ILLUMINA | Illumina HiSeq 2000 | PAIRED |
| **ERR479325** | ERS436659 | ERP005534 | France | Europe | 0 | ILLUMINA | Illumina HiSeq 2000 | PAIRED |
| **ERR479326** | ERS436659 | ERP005534 | France | Europe | 0 | ILLUMINA | Illumina HiSeq 2000 | PAIRED |
| **ERR479063** | ERS436660 | ERP005534 | France | Europe | 0 | ILLUMINA | Illumina HiSeq 2000 | PAIRED |
| **ERR479064** | ERS436660 | ERP005534 | France | Europe | 0 | ILLUMINA | Illumina HiSeq 2000 | PAIRED |
| **ERR479099** | ERS436664 | ERP005534 | France | Europe | 0 | ILLUMINA | Illumina HiSeq 2000 | PAIRED |
| **ERR479100** | ERS436664 | ERP005534 | France | Europe | 0 | ILLUMINA | Illumina HiSeq 2000 | PAIRED |
| **ERR479274** | ERS436666 | ERP005534 | France | Europe | 0 | ILLUMINA | Illumina HiSeq 2000 | PAIRED |
| **ERR479275** | ERS436666 | ERP005534 | France | Europe | 0 | ILLUMINA | Illumina HiSeq 2000 | PAIRED |
| **ERR479004** | ERS436669 | ERP005534 | France | Europe | 0 | ILLUMINA | Illumina HiSeq 2000 | PAIRED |

| | | | | | | | | |
|---|---|---|---|---|---|---|---|---|
| **ERR479005** | ERS436669 | ERP005534 | France | Europe | 0 | ILLUMINA | Illumina HiSeq 2000 | PAIRED |
| **ERR479043** | ERS436718 | ERP005534 | France | Europe | 0 | ILLUMINA | Illumina HiSeq 2000 | PAIRED |
| **ERR479044** | ERS436718 | ERP005534 | France | Europe | 0 | ILLUMINA | Illumina HiSeq 2000 | PAIRED |
| **ERR478964** | ERS436722 | ERP005534 | France | Europe | 0 | ILLUMINA | Illumina HiSeq 2000 | PAIRED |
| **ERR478965** | ERS436722 | ERP005534 | France | Europe | 0 | ILLUMINA | Illumina HiSeq 2000 | PAIRED |
| **ERR478966** | ERS436722 | ERP005534 | France | Europe | 0 | ILLUMINA | Illumina HiSeq 2000 | PAIRED |
| **ERR478967** | ERS436722 | ERP005534 | France | Europe | 0 | ILLUMINA | Illumina HiSeq 2000 | PAIRED |
| **ERR479321** | ERS436738 | ERP005534 | France | Europe | 0 | ILLUMINA | Illumina HiSeq 2000 | PAIRED |
| **ERR479322** | ERS436738 | ERP005534 | France | Europe | 0 | ILLUMINA | Illumina HiSeq 2000 | PAIRED |
| **ERR479323** | ERS436738 | ERP005534 | France | Europe | 0 | ILLUMINA | Illumina HiSeq 2000 | PAIRED |
| **ERR479324** | ERS436738 | ERP005534 | France | Europe | 0 | ILLUMINA | Illumina HiSeq 2000 | PAIRED |
| **ERR479267** | ERS436739 | ERP005534 | France | Europe | 0 | ILLUMINA | Illumina HiSeq 2000 | PAIRED |
| **ERR479268** | ERS436739 | ERP005534 | France | Europe | 0 | ILLUMINA | Illumina HiSeq 2000 | PAIRED |
| **ERR479269** | ERS436739 | ERP005534 | France | Europe | 0 | ILLUMINA | Illumina HiSeq 2000 | PAIRED |
| **ERR479341** | ERS436742 | ERP005534 | France | Europe | 0 | ILLUMINA | Illumina HiSeq 2000 | PAIRED |
| **ERR479342** | ERS436742 | ERP005534 | France | Europe | 0 | ILLUMINA | Illumina HiSeq 2000 | PAIRED |
| **ERR479022** | ERS436767 | ERP005534 | France | Europe | 0 | ILLUMINA | Illumina HiSeq 2000 | PAIRED |
| **ERR479023** | ERS436767 | ERP005534 | France | Europe | 0 | ILLUMINA | Illumina HiSeq 2000 | PAIRED |
| **ERR478994** | ERS436771 | ERP005534 | France | Europe | 0 | ILLUMINA | Illumina HiSeq 2000 | PAIRED |
| **ERR478995** | ERS436771 | ERP005534 | France | Europe | 0 | ILLUMINA | Illumina HiSeq 2000 | PAIRED |
| **ERR478996** | ERS436771 | ERP005534 | France | Europe | 0 | ILLUMINA | Illumina HiSeq 2000 | PAIRED |
| **ERR478997** | ERS436771 | ERP005534 | France | Europe | 0 | ILLUMINA | Illumina HiSeq 2000 | PAIRED |
| **ERR479118** | ERS436772 | ERP005534 | France | Europe | 0 | ILLUMINA | Illumina HiSeq 2000 | PAIRED |
| **ERR479119** | ERS436772 | ERP005534 | France | Europe | 0 | ILLUMINA | Illumina HiSeq 2000 | PAIRED |
| **ERR479120** | ERS436772 | ERP005534 | France | Europe | 0 | ILLUMINA | Illumina HiSeq 2000 | PAIRED |
| **ERR479121** | ERS436772 | ERP005534 | France | Europe | 0 | ILLUMINA | Illumina HiSeq 2000 | PAIRED |
| **ERR478986** | ERS436785 | ERP005534 | France | Europe | 0 | ILLUMINA | Illumina HiSeq 2000 | PAIRED |
| **ERR478987** | ERS436785 | ERP005534 | France | Europe | 0 | ILLUMINA | Illumina HiSeq 2000 | PAIRED |

| | | | | | | | | |
|---|---|---|---|---|---|---|---|---|
| **ERR478988** | ERS436785 | ERP005534 | France | Europe | 0 | ILLUMINA | Illumina HiSeq 2000 | PAIRED |
| **ERR478989** | ERS436785 | ERP005534 | France | Europe | 0 | ILLUMINA | Illumina HiSeq 2000 | PAIRED |
| **ERR478976** | ERS436788 | ERP005534 | France | Europe | 0 | ILLUMINA | Illumina HiSeq 2000 | PAIRED |
| **ERR478977** | ERS436788 | ERP005534 | France | Europe | 0 | ILLUMINA | Illumina HiSeq 2000 | PAIRED |
| **ERR478978** | ERS436788 | ERP005534 | France | Europe | 0 | ILLUMINA | Illumina HiSeq 2000 | PAIRED |
| **ERR478979** | ERS436788 | ERP005534 | France | Europe | 0 | ILLUMINA | Illumina HiSeq 2000 | PAIRED |
| **ERR479209** | ERS436791 | ERP005534 | France | Europe | 0 | ILLUMINA | Illumina HiSeq 2000 | PAIRED |
| **ERR479210** | ERS436791 | ERP005534 | France | Europe | 0 | ILLUMINA | Illumina HiSeq 2000 | PAIRED |
| **ERR479211** | ERS436791 | ERP005534 | France | Europe | 0 | ILLUMINA | Illumina HiSeq 2000 | PAIRED |
| **ERR479212** | ERS436791 | ERP005534 | France | Europe | 0 | ILLUMINA | Illumina HiSeq 2000 | PAIRED |
| **ERR479570** | ERS436839 | ERP005534 | Germany | Europe | 0 | ILLUMINA | Illumina HiSeq 2000 | PAIRED |
| **ERR479571** | ERS436839 | ERP005534 | Germany | Europe | 0 | ILLUMINA | Illumina HiSeq 2000 | PAIRED |
| **ERR479572** | ERS436839 | ERP005534 | Germany | Europe | 0 | ILLUMINA | Illumina HiSeq 2000 | PAIRED |
| **ERR479573** | ERS436839 | ERP005534 | Germany | Europe | 0 | ILLUMINA | Illumina HiSeq 2000 | PAIRED |
| **ERR479574** | ERS436839 | ERP005534 | Germany | Europe | 0 | ILLUMINA | Illumina HiSeq 2000 | PAIRED |
| **ERR479575** | ERS436839 | ERP005534 | Germany | Europe | 0 | ILLUMINA | Illumina HiSeq 2000 | PAIRED |
| **ERR479590** | ERS436841 | ERP005534 | Germany | Europe | 0 | ILLUMINA | Illumina HiSeq 2000 | PAIRED |
| **ERR479591** | ERS436841 | ERP005534 | Germany | Europe | 0 | ILLUMINA | Illumina HiSeq 2000 | PAIRED |
| **ERR479592** | ERS436841 | ERP005534 | Germany | Europe | 0 | ILLUMINA | Illumina HiSeq 2000 | PAIRED |
| **ERR479593** | ERS436841 | ERP005534 | Germany | Europe | 0 | ILLUMINA | Illumina HiSeq 2000 | PAIRED |
| **ERR479594** | ERS436841 | ERP005534 | Germany | Europe | 0 | ILLUMINA | Illumina HiSeq 2000 | PAIRED |
| **ERR479595** | ERS436841 | ERP005534 | Germany | Europe | 0 | ILLUMINA | Illumina HiSeq 2000 | PAIRED |
| **ERR479596** | ERS436841 | ERP005534 | Germany | Europe | 0 | ILLUMINA | Illumina HiSeq 2000 | PAIRED |
| **SRR19064371** | SRS12879096 | SRP373424 | United States of America | North America | 0 | ILLUMINA | Illumina NovaSeq 6000 | PAIRED |
| **SRR19064376** | SRS12879092 | SRP373424 | United States of America | North America | 0 | ILLUMINA | Illumina NovaSeq 6000 | PAIRED |
| **SRR19064708** | SRS12879194 | SRP373424 | United States of America | North America | 0 | ILLUMINA | Illumina NovaSeq 6000 | PAIRED |
| **SRR19064969** | SRS12879374 | SRP373424 | United States of America | North America | 0 | ILLUMINA | Illumina NovaSeq 6000 | PAIRED |
| **SRR19064372** | SRS12879094 | SRP373424 | United States of America | North America | 0 | ILLUMINA | Illumina NovaSeq 6000 | PAIRED |

| | | | | | | | | |
|---|---|---|---|---|---|---|---|---|
| **SRR19064377** | SRS12879089 | SRP373424 | United States of America | North America | 0 | ILLUMINA | Illumina NovaSeq 6000 | PAIRED |
| **SRR19064416** | SRS12879050 | SRP373424 | United States of America | North America | 0 | ILLUMINA | Illumina NovaSeq 6000 | PAIRED |
| **SRR19064446** | SRS12879020 | SRP373424 | United States of America | North America | 0 | ILLUMINA | Illumina NovaSeq 6000 | PAIRED |
| **SRR19064643** | SRS12879259 | SRP373424 | United States of America | North America | 0 | ILLUMINA | Illumina NovaSeq 6000 | PAIRED |
| **SRR19064650** | SRS12879252 | SRP373424 | United States of America | North America | 0 | ILLUMINA | Illumina NovaSeq 6000 | PAIRED |
| **SRR19064695** | SRS12879206 | SRP373424 | United States of America | North America | 0 | ILLUMINA | Illumina NovaSeq 6000 | PAIRED |
| **SRR19064741** | SRS12879160 | SRP373424 | United States of America | North America | 0 | ILLUMINA | Illumina NovaSeq 6000 | PAIRED |
| **SRR19064901** | SRS12879442 | SRP373424 | United States of America | North America | 0 | ILLUMINA | Illumina NovaSeq 6000 | PAIRED |
| **SRR19064921** | SRS12879422 | SRP373424 | United States of America | North America | 0 | ILLUMINA | Illumina NovaSeq 6000 | PAIRED |
| **SRR19064389** | SRS12879081 | SRP373424 | United States of America | North America | 0 | ILLUMINA | Illumina NovaSeq 6000 | PAIRED |
| **SRR19064451** | SRS12879016 | SRP373424 | United States of America | North America | 0 | ILLUMINA | Illumina NovaSeq 6000 | PAIRED |
| **SRR19064600** | SRS12879302 | SRP373424 | United States of America | North America | 0 | ILLUMINA | Illumina NovaSeq 6000 | PAIRED |
| **SRR19064677** | SRS12879226 | SRP373424 | United States of America | North America | 0 | ILLUMINA | Illumina NovaSeq 6000 | PAIRED |
| **SRR19064715** | SRS12879187 | SRP373424 | United States of America | North America | 0 | ILLUMINA | Illumina NovaSeq 6000 | PAIRED |
| **SRR19064809** | SRS12879532 | SRP373424 | United States of America | North America | 0 | ILLUMINA | Illumina NovaSeq 6000 | PAIRED |

**Table S8. 10-strain *Escherichia coli* relative abundances mix for the real-world metagenomes multi-strain spike-ins dataset.** 10 *E. coli* strains were spiked in at 1% total read count into the 42 metagenome samples listed in Table S7. 5 intra-phylogroup B2-2 strains (PMV-1, IHE3034, SF-468, D8 and STEC299) were spiked in at equal relative abundance (0.1), and 5 broader phylogroup diversity strains were spiked in at differential relative abundance (0.04–0.20).

| ***E. coli* strain** | **Accession ID** | **Phylogroup** | **Relative abundance** |
|---|---|---|---|
| PMV-1 | GCA_000493595.1_EcoPMV1_genomic | B2-2 | 0.1 |
| IHE3034 | GCA_000025745.1_ASM2574v1_genomic | B2-2 | 0.1 |
| SF-468 | GCA_001280345.1_ASM128034v1_genomic | B2-2 | 0.1 |
| D8 | GCA_001900395.1_ASM190039v1_genomic | B2-2 | 0.1 |
| STEC299 | GCA_002863845.1_ASM286384v1_genomic | B2-2 | 0.1 |
| AR_0081 | GCA_002996945.1_ASM299694v1_genomic | B2-1 | 0.2 |
| SC516 | GCA_002812545.1_ASM281254v1_genomic | A | 0.12 |
| AR437 | GCA_003073815.1_ASM307381v1_genomic | B1 | 0.08 |
| RM12579 | GCA_000245515.1_ASM24551v1_genomic | E2 | 0.06 |
| AR_0077 | GCA_002948655.1_ASM294865v1_genomic | F | 0.04 |

**Table S9. Genomic relatedness of PathoScope and StrainGE predictions to removed ground-truth strains.** The 10 ground-truth *E. coli* strains were removed from the 304-strain reference database prior to strain-level profiling of the 42 metagenomes from the real-world spike-ins dataset. For each removed ground-truth strain, the detected same-phylogroup reference by PathoScope and StrainGE with the highest pairwise ANI is shown. "Samples detected" indicates the number of metagenomes ($n$ = 42) in which that specific reference strain was detected following application of the 0.005 relative-abundance cutoff. Pairwise ANI represents the genomic similarity between the removed ground-truth strain and the predicted strain. ANI rank indicates the rank of the predicted strain among the remaining 294 available reference genomes, where rank 1 represents the closest available reference by ANI.

| Tool | Removed ground-truth strain | Phylogroup | Closest predicted strain | Samples detected in ($n$ = 42) | Predicted phylogroup | Pairwise ANI (%) | ANI rank among 294 references |
|---|---|---|---|---|---|---|---|
| StrainGE | SC516 | A | S30 | 41 | A | 99.0507 | 1 |
| PathoScope | SC516 | A | S30 | 24 | A | 99.0507 | 1 |
| StrainGE | AR437 | B1 | FDAARGOS_433 | 42 | B1 | 99.9034 | 1 |
| PathoScope | AR437 | B1 | FDAARGOS_433 | 42 | B1 | 99.9034 | 1 |
| StrainGE | AR_0081 | B2-1 | O25b:H4 | 42 | B2-1 | 99.828 | 4 |
| PathoScope | AR_0081 | B2-1 | MNCRE44 | 4 | B2-1 | 99.8511 | 1 |
| StrainGE | IHE3034 | B2-2 | APEC O18 | 42 | B2-2 | 99.9048 | 1 |
| PathoScope | IHE3034 | B2-2 | APEC O18 | 42 | B2-2 | 99.9048 | 1 |
| StrainGE | PMV-1 | B2-2 | APEC O18 | 42 | B2-2 | 99.8766 | 1 |
| PathoScope | PMV-1 | B2-2 | APEC O18 | 42 | B2-2 | 99.8766 | 1 |
| StrainGE | SF-468 | B2-2 | APEC O18 | 42 | B2-2 | 99.6727 | 4 |
| PathoScope | SF-468 | B2-2 | HS13-1 | 36 | B2-2 | 99.7716 | 1 |
| StrainGE | D8 | B2-2 | APEC O18 | 42 | B2-2 | 99.091 | 1 |
| PathoScope | D8 | B2-2 | APEC O18 | 42 | B2-2 | 99.091 | 1 |
| StrainGE | STEC299 | B2-2 | APEC O18 | 42 | B2-2 | 98.8585 | 5 |
| PathoScope | STEC299 | B2-2 | APEC O18 | 42 | B2-2 | 98.8585 | 5 |
| StrainGE | RM12579 | E2 | 2013C-4465 | 42 | E2 | 99.8876 | 1 |
| PathoScope | RM12579 | E2 | 2013C-4465 | 42 | E2 | 99.8876 | 1 |
| StrainGE | AR_0077 | F | N/A | 0 | N/A | N/A | N/A |
| PathoScope | AR_0077 | F | SMS-3-5 | 42 | F | 98.4675 | 6 |

<u>Supplementary Figures</u>

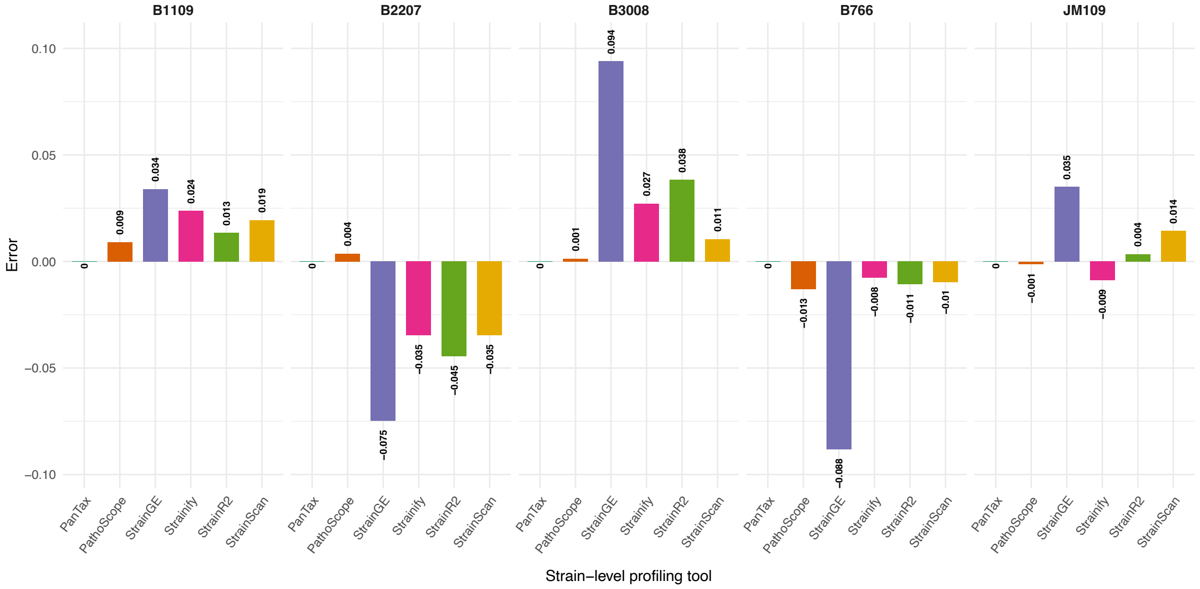


**Figure S1. Error of five equally abundant *E. coli* strains across strain-level profiling tools in the ZymoBIOMICS® D6331 gut microbiome standard**. All tools were run on the ZymoBIOMICS® standard with the five target strains included in the reference database, and predicted relative abundances were used to calculate error for each tool. Faceted by strain to show the over- and underestimation of relative abundance predictions relative to the ground truth (0.2; Table S2).

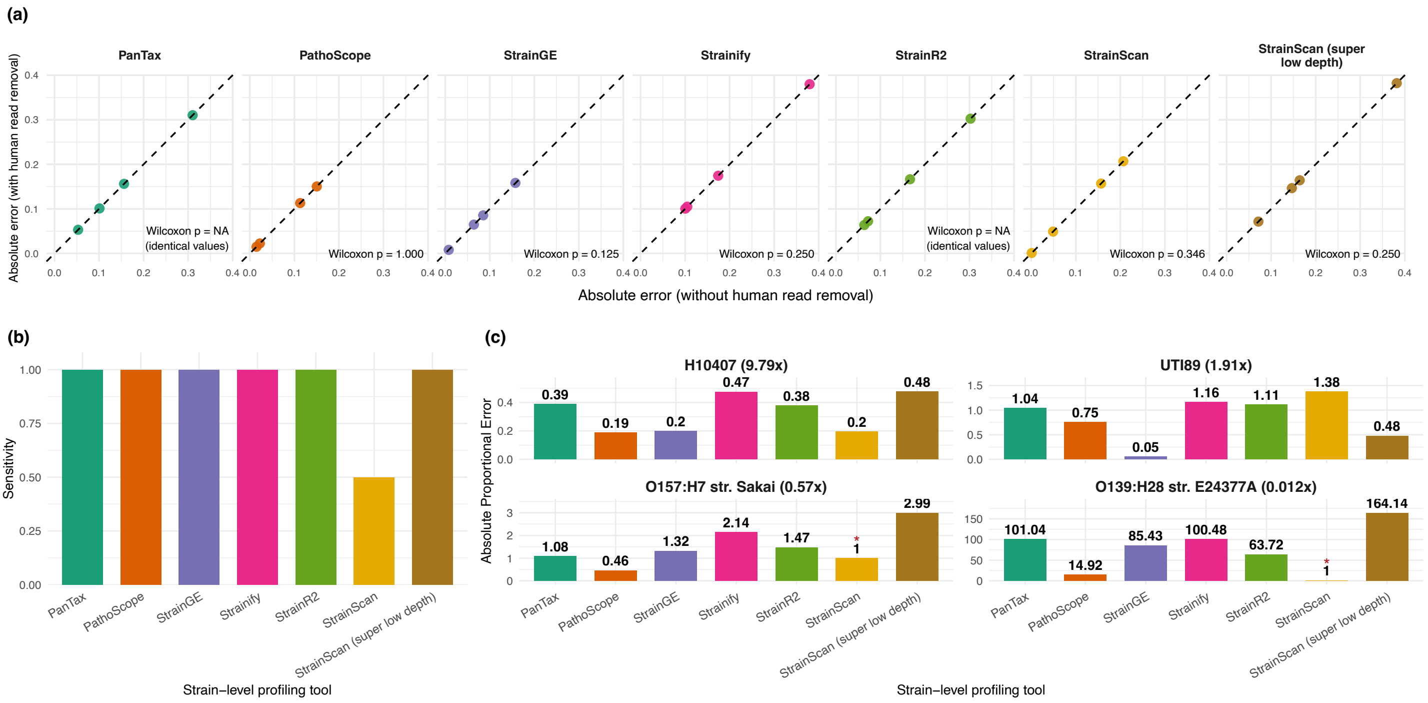


**Figure S2. Prediction error with and without human read removal, sensitivity and absolute proportional error across strain-level profiling tools in the SRR13355226 mock community.** (a) Paired Wilcoxon signed-rank tests on absolute error per tool, comparing conditions with and without human read removal (Deacon). A significance threshold of $P < 0.05$ was applied. No significant differences were observed. (b) Sensitivity per base tool, collapsed per base tool and calculated from true positive and false negative call-type counts (see Methods). (c) Absolute proportional error per base tool across four differentially abundant *E. coli* strains. Facets indicate strain and estimated sequencing depth (x). An asterisk (*) denotes that strain not detected, for which an APE of 1 was assigned (see Methods). Note that *y*-axis scales differ between facets.

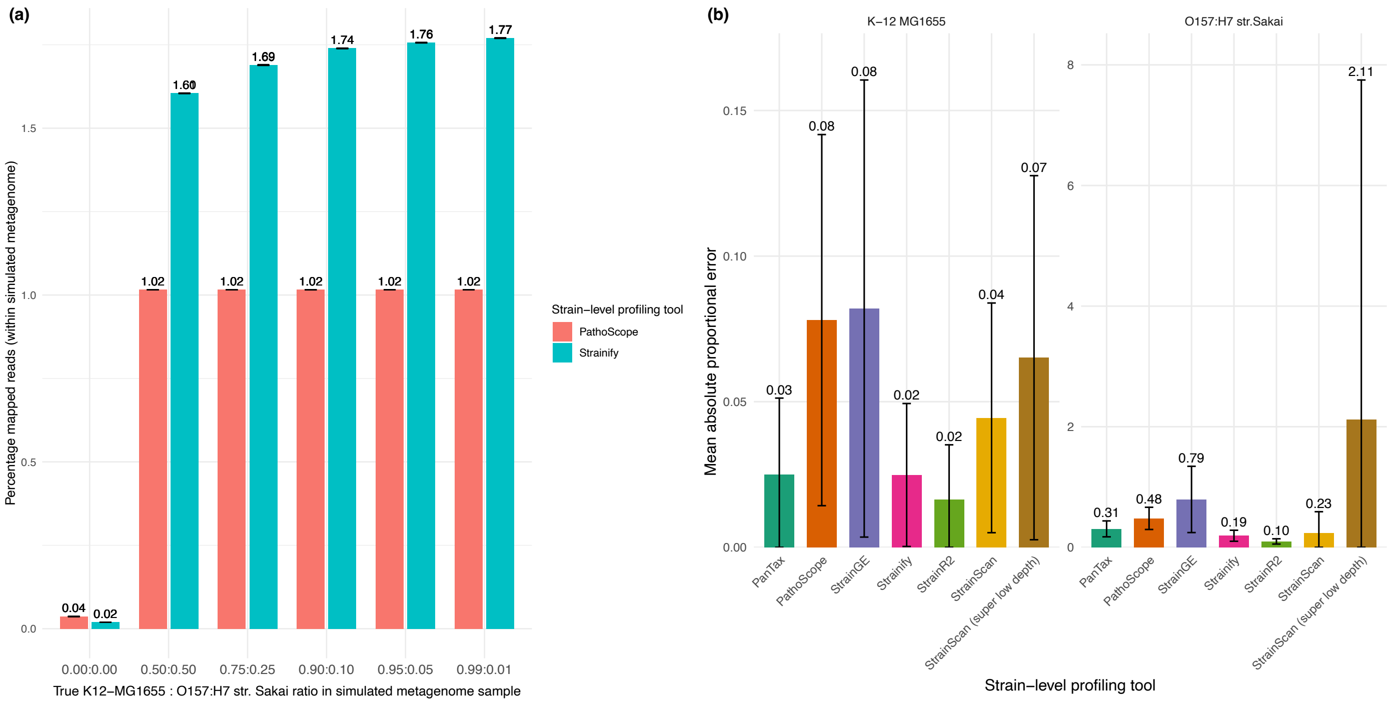


**Figure S3. Percentage of reads mapped and mean absolute proportional error per true *E. coli* strain (K12-MG1655 and O157:H7 Sakai) in the simulated metagenome dataset.** (a) Mean percentage of mapped reads per sample for PathoScope and Strainify, averaged across technical triplicates and all sequencing efforts (20, 50 and 100M showed no difference in values to two decimal places (see Data Summary)). (b) Mean absolute proportional error of predicted relative abundances for each true strain (K12-MG1655 and O157:H7 str. Sakai) across tools. Error bars represent mean ± standard deviation. Note that different y-axis scales are used for each strain.

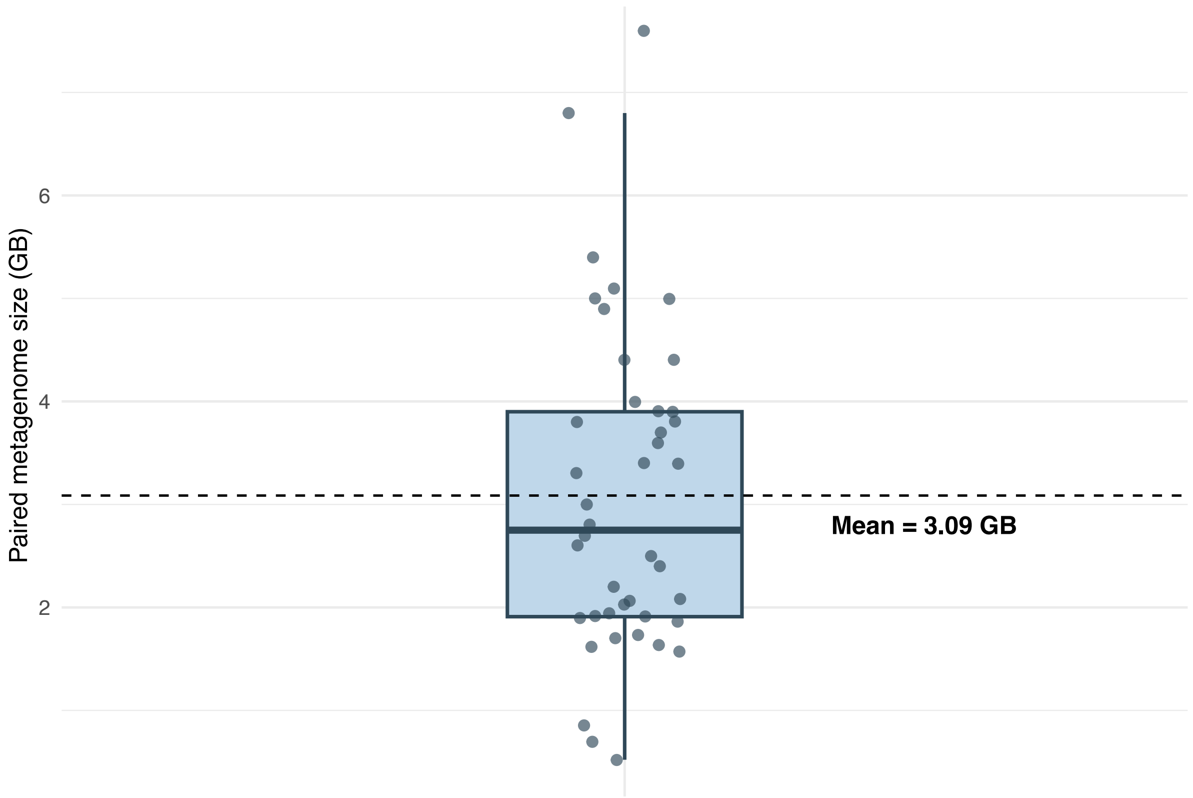


**Figure S4. Paired metagenome file sizes comprising the real-world multi-strain spike-in dataset.** Points represent individual metagenome samples, with file size calculated as the combined R1 + R2 size following quality control using metagen-fetch (https://github.com/alexmsalmeida/metagen-fetch). Paired metgenome file size ranged from 0.52–7.60 GB.

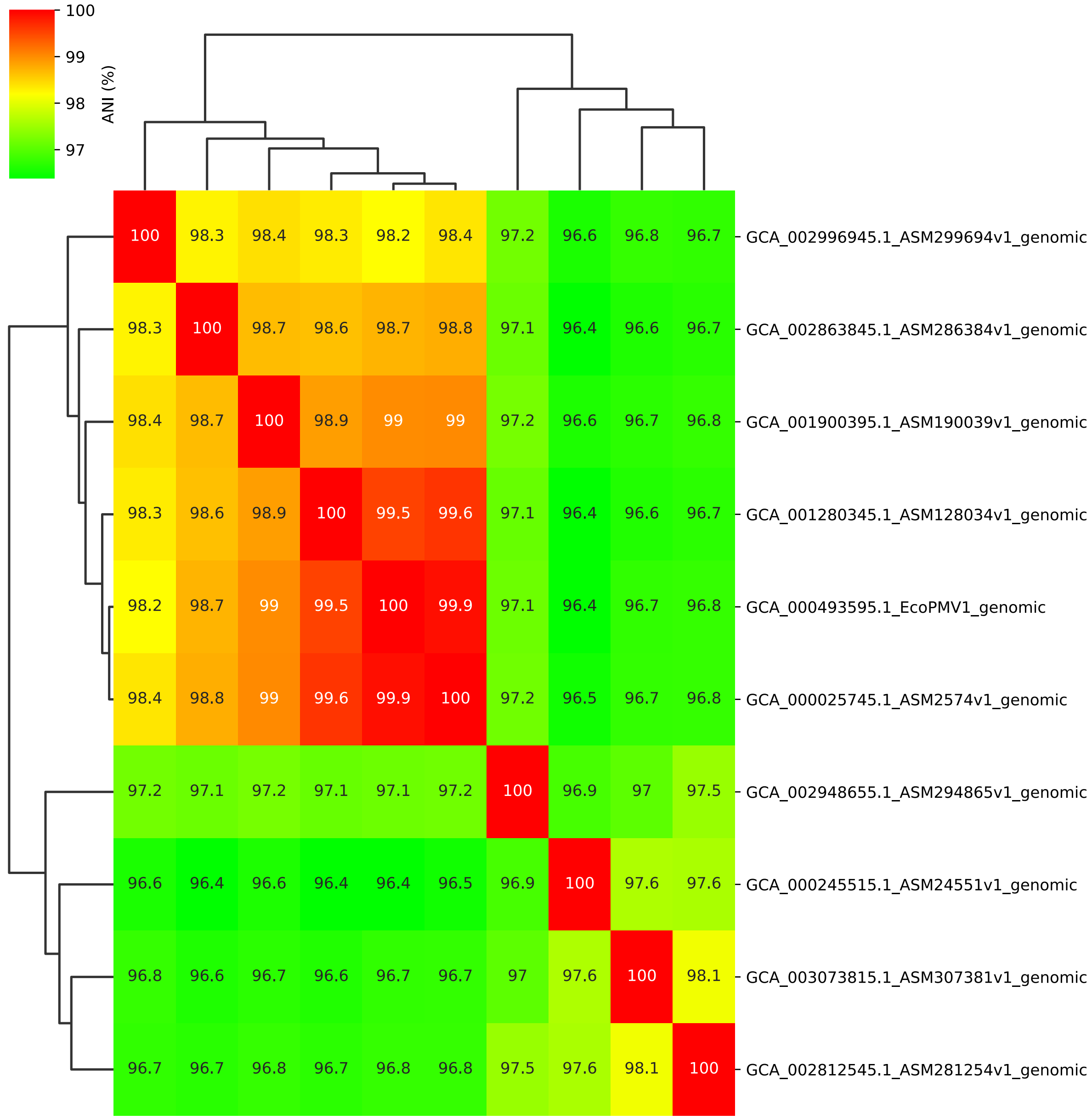


**Fig. S5. Pairwise whole-genome ANI among the 10 E. coli strains used in the real-world spike-ins dataset.** Heatmap values show all-vs-all ANI (%) calculated using FastANI v1.34[17], with hierarchical clustering indicating genomic relatedness. Strain metadata are provided in Table S8.

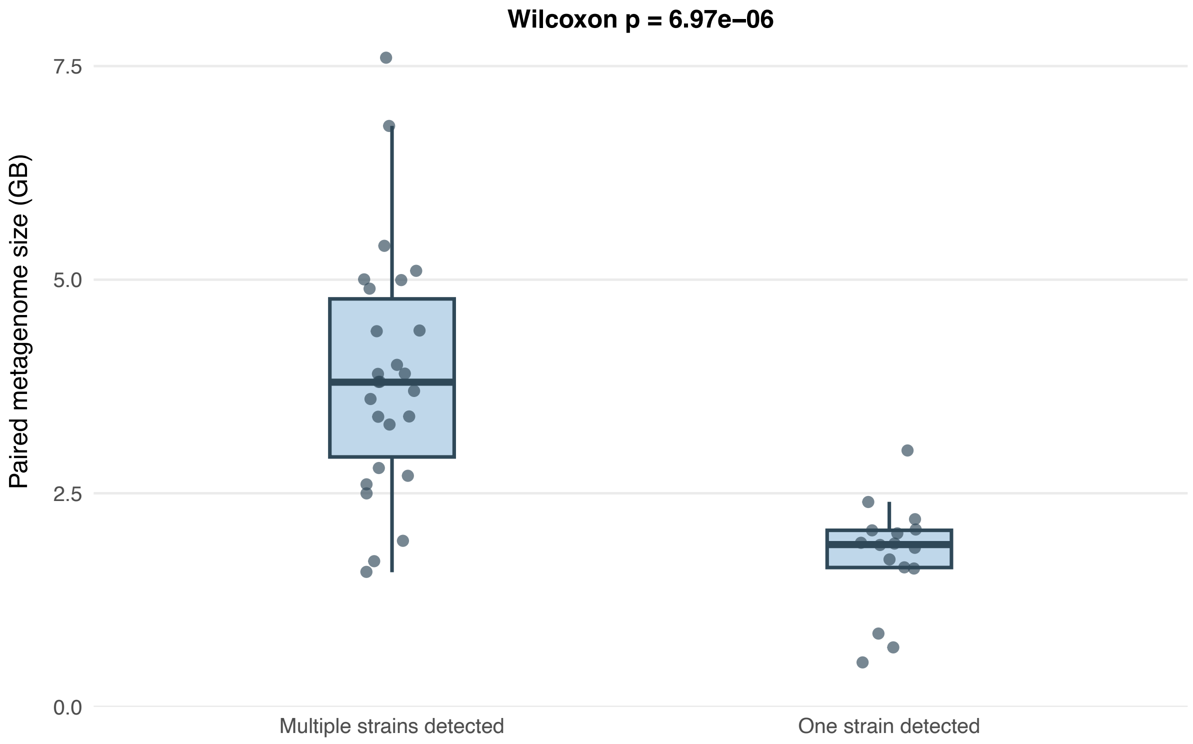


**Fig. S6. Association between StrainScan strain detection and metagenome size in the real-world multi-strain spike-in dataset.** Paired metagenome file size (R1 + R2; GB) was compared between samples in which StrainScan detected one strain versus multiple strains. Points represent individual metagenome samples. Samples with only one detected strain had significantly smaller metagenome sizes than those with multiple detected strains (Wilcoxon rank-sum test, $p = 6.97 \times 10^{-6}$).

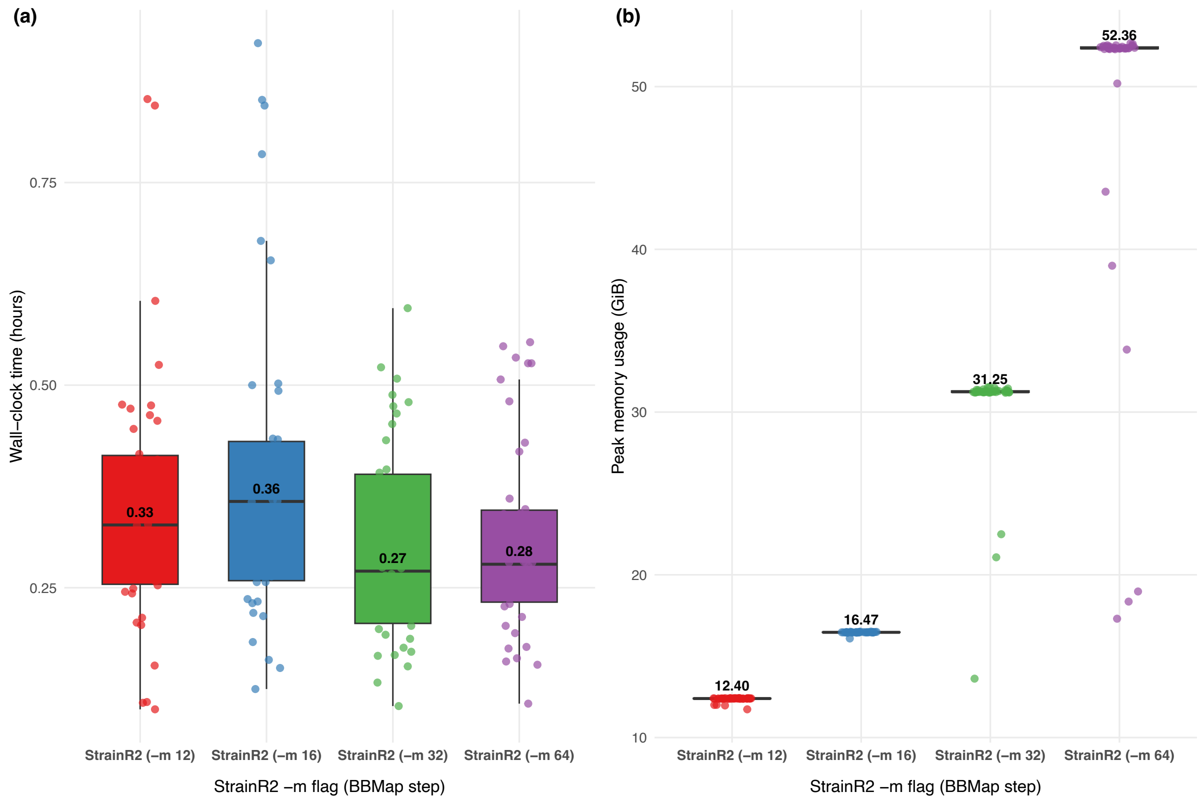


**Fig. S7. Effect of the StrainR2 memory allocation (-m) parameter on metagenome profiling runtime and peak memory usage.** Wall-clock runtime (a) and peak memory usage (b) were recorded across the 42 real-world spike-ins dataset metagenomes for StrainR2 -m settings of 12, 16, 32 and 64 GB. The -m parameter controls the gigabytes of memory to use when running the short-read aligner BBMap[53]. Points represent individual metagenome runs, box plots show the median and interquartile range, and values above the box plots indicate medians. Results were identical across -m parameters (data not shown). Computational resource usage was recorded using */usr/bin/time -v* command line utility. An -m setting of 8 GiB was also tested but failed across all metagenome samples and is therefore not shown.

### References (Supplementary Material ONLY):


1. Zhang W, Liu Y, Li G, Xu J, Chen E, Schönhuth A, et al. Strain-level metagenomic profiling using pangenome graphs with PanTax. Genome Res. 2026. doi: 10.1101/gr.280858.125
2. Gurobi Optimization L. Gurobi Optimizer Reference Manual. 11.0 ed2026.
3. Chang X, Novak AM, Eizenga JM, Sirén J, Monlong J, Negi S, et al. Rapid, accurate long- and short-read mapping to large pangenome graphs with vg Giraffe. bioRxiv. 2025. doi: 10.1101/2025.09.29.678807
4. Hong C, Manimaran S, Shen Y, Perez-Rogers JF, Byrd AL, Castro-Nallar E, et al. PathoScope 2.0: a complete computational framework for strain identification in environmental or clinical sequencing samples. Microbiome. 2014;2:33. doi: 10.1186/2049-2618-2-33
5. van Dijk LR, Walker BJ, Straub TJ, Worby CJ, Grote A, Schreiber HLt, et al. StrainGE: a toolkit to track and characterize low-abundance strains in complex microbial communities. Genome Biol. 2022;23(1):74. doi: 10.1186/s13059-022-02630-0
6. Luo RS, Kille B, Vaughan EE, Clark JR, Maresso AW, Nute MG, et al. Strainify: Strain-Level Microbiome Profiling for Low-Coverage Short-Read Metagenomic Datasets. bioRxiv. 2025. doi: 10.1101/2025.10.10.681738
7. Heber K, Tian S, Betancurt-Anzola D, Koo H, Bisanz JE. StrainR2 accurately deconvolutes strain-level abundances in synthetic microbial communities. Bioinformatics. 2025;41(8). doi: 10.1093/bioinformatics/btaf440
8. Liao H, Ji Y, Sun Y. High-resolution strain-level microbiome composition analysis from short reads. Microbiome. 2023;11(1):183. doi: 10.1186/s40168-023-01615-w
9. Waters NR, Abram F, Brennan F, Holmes A, Pritchard L. Easy phylotyping of Escherichia coli via the EzClermont web app and command-line tool. Access Microbiol. 2020;2(9):acmi000143. doi: 10.1099/acmi.0.000143
10. Seemann T. mlst: Github; [Available from: https://github.com/tseemann/mlst.
11. Jolley KA, Bray JE, Maiden MCJ. Open-access bacterial population genomics: BIGSdb software, the PubMLST.org website and their applications. Wellcome Open Res. 2018;3:124. doi: 10.12688/wellcomeopenres.14826.1
12. Wenger AM, Peluso P, Rowell WJ, Chang PC, Hall RJ, Concepcion GT, et al. Accurate circular consensus long-read sequencing improves variant detection and assembly of a human genome. Nat Biotechnol. 2019;37(10):1155–62. doi: 10.1038/s41587-019-0217-9
13. Constantinides B, Lees J, Crook DW. Deacon: fast sequence filtering and contaminant depletion. bioRxiv. 2025:2025.06.09.658732. doi: 10.1101/2025.06.09.658732
14. King CH, Desai H, Sylvetsky AC, LoTempio J, Ayanyan S, Carrie J, et al. Baseline human gut microbiota profile in healthy people and standard reporting template. PLoS One. 2019;14(9):e0206484. doi: 10.1371/journal.pone.0206484
15. Gourlé H, Karlsson-Lindsjö O, Hayer J, Bongcam-Rudloff E. Simulating Illumina metagenomic data with InSilicoSeq. Bioinformatics. 2019;35(3):521–2. doi: 10.1093/bioinformatics/bty630
16. Abram K, Udaondo Z, Bleker C, Wanchai V, Wassenaar TM, Robeson MS, 2nd, et al. Mash-based analyses of Escherichia coli genomes reveal 14 distinct phylogroups. Commun Biol. 2021;4(1):117. doi: 10.1038/s42003-020-01626-5
17. Jain C, Rodriguez RL, Phillippy AM, Konstantinidis KT, Aluru S. High throughput ANI analysis of 90K prokaryotic genomes reveals clear species boundaries. Nat Commun. 2018;9(1):5114. doi: 10.1038/s41467-018-07641-9
18. Yin Q, da Silva AC, Zorrilla F, Almeida AS, Patil KR, Almeida A. Ecological dynamics of Enterobacteriaceae in the human gut microbiome across global populations. Nat Microbiol. 2025;10(2):541–53. doi: 10.1038/s41564-024-01912-6
19. Wood DE, Lu J, Langmead B. Improved metagenomic analysis with Kraken 2. Genome Biol. 2019;20(1):257. doi: 10.1186/s13059-019-1891-0

20. Lu J, Rincon N, Wood DE, Breitwieser FP, Pockrandt C, Langmead B, et al. Metagenome analysis using the Kraken software suite. Nat Protoc. 2022;17(12):2815–39. doi: 10.1038/s41596-022-00738-y
21. O'Leary NA, Cox E, Holmes JB, Anderson WR, Falk R, Hem V, et al. Exploring and retrieving sequence and metadata for species across the tree of life with NCBI Datasets. Sci Data. 2024;11(1):732. doi: 10.1038/s41597-024-03571-y
22. Olm MR, Brown CT, Brooks B, Banfield JF. dRep: a tool for fast and accurate genomic comparisons that enables improved genome recovery from metagenomes through de-replication. Isme j. 2017;11(12):2864–8. doi: 10.1038/ismej.2017.126
23. R Core Team. R: A Language and Environment for Statistical Computing. 4.5.1 ed. Vienna, Austria: R Foundation for Statistical Computing; 2025.
24. Wickham H, Bryan J. readxl: Read Excel Files (R package version 1.4.5). 2025.
25. Wickham H. ggplot2: Elegant Graphics for Data Analysis. Springer-Verlag New York. 2016. doi: 10.1007/978-3-319-24277-4_9
26. Wilke CO. cowplot: Streamlined Plot Theme and Plot Annotations for “ggplot2” (R package version 1.2.0). 2024.
27. Wickham H, François R, Henry L, Müller K, Vaughan D. dplyr: A Grammar of Data Manipulation. 2023.
28. Wickham H. stringr: Simple, Consistent Wrappers for Common String Operations (R package version 1.6.0). 2025.
29. Wickham H, Vaughan A, Girlich M. tidyr: Tidy Messy Data (R package version 1.3.1). 2024.
30. Wickham H. forcats: Tools for Working with Categorical Variables (Factors; R package version 1.0.1). 2025.
31. Wickham H, Pedersen T, Seidel D. Scales: Scale Functions for Visualization (R package version 1.4.0). 2025.
32. Dorai-Raj S. binom: Binomial Confidence Intervals for Several Parameterizations (R package version 1.1.1.1). 2022.
33. Wickham H, Henry L. purrr: Functional Programming Tools (R package version 1.1.0). 2025.
34. Xu S, Chen M, Feng T, Zhan L, Zhou L, Yu G. Use ggbreak to Effectively Utilize Plotting Space to Deal With Large Datasets and Outliers. (R package version 0.1.7). Front Genet. 2021;12:774846. doi: 10.3389/fgene.2021.774846
35. Neuwirth E. RColorBrewer: ColorBrewer Palettes. (R package version 1.1.3). 2022.
36. Zeileis A, Fisher JC, Hornik K, Ihaka R, McWhite CD, Murrell P, et al. colorspace: A Toolbox for Manipulating and Assessing Colors and Palettes. (R package version 2.1.2). Journal of Statistical Software. 2020;96(1):1 – 49. doi: 10.18637/jss.v096.i01
37. Wilke C, Wiernik B. ggtext: Improved Text Rendering Support for 'ggplot2'. (R package version 0.2.0). 2026.
38. Yu G. ggplotify: Convert Plot to 'grob' or 'ggplot' Object. (R package version 0.1.3). 2025.
39. Albanese D, Donati C. Strain profiling and epidemiology of bacterial species from metagenomic sequencing. Nat Commun. 2017;8(1):2260. doi: 10.1038/s41467-017-02209-5
40. Da Silva K, Pons N, Berland M, Plaza Oñate F, Almeida M, Peterlongo P. StrainFLAIR: strain-level profiling of metagenomic samples using variation graphs. PeerJ. 2021;9:e11884. doi: 10.7717/peerj.11884
41. Li X, Hu H, Li X. mixtureS: a novel tool for bacterial strain genome reconstruction from reads. Bioinformatics. 2021;37(4):575–7. doi: 10.1093/bioinformatics/btaa728
42. France M, Chaudry I, Elsherbini J, Ravel J. kSanity: A k-mer based application for precision bacterial strain detection and quantification. bioRxiv. 2025. doi: 10.1101/2025.09.04.674052
43. Blanco-Míguez A, Beghini F, Cumbo F, McIver LJ, Thompson KN, Zolfo M, et al. Extending and improving metagenomic taxonomic profiling with uncharacterized species using MetaPhlAn 4. Nat Biotechnol. 2023;41(11):1633–44. doi: 10.1038/s41587-023-01688-w

44. Murovec B, Deutsch L, Osredkar D, Stres B. MetaBakery: a Singularity implementation of bioBakery tools as a skeleton application for efficient HPC deconvolution of microbiome metagenomic sequencing data to machine learning ready information. Front Microbiol. 2024;15:1426465. doi: 10.3389/fmicb.2024.1426465
45. Beghini F, McIver LJ, Blanco-Míguez A, Dubois L, Asnicar F, Maharjan S, et al. Integrating taxonomic, functional, and strain-level profiling of diverse microbial communities with bioBakery 3. Elife. 2021;10. doi: 10.7554/eLife.65088
46. Ahn TH, Chai J, Pan C. Sigma: strain-level inference of genomes from metagenomic analysis for biosurveillance. Bioinformatics. 2015;31(2):170–7. doi: 10.1093/bioinformatics/btu641
47. Luo C, Knight R, Siljander H, Knip M, Xavier RJ, Gevers D. ConStrains identifies microbial strains in metagenomic datasets. Nat Biotechnol. 2015;33(10):1045–52. doi: 10.1038/nbt.3319
48. Lin H-N, Lin Y-L, Hsu W-L. StrainPro – a highly accurate Metagenomic strain-level profiling tool. bioRxiv. 2019:807149. doi: 10.1101/807149
49. Shaw J, Gounot JS, Chen H, Nagarajan N, Yu YW. Floria: fast and accurate strain haplotyping in metagenomes. Bioinformatics. 2024;40(Suppl 1):i30–i8. doi: 10.1093/bioinformatics/btae252
50. Quince C, Nurk S, Raguideau S, James R, Soyer OS, Summers JK, et al. STRONG: metagenomics strain resolution on assembly graphs. Genome Biol. 2021;22(1):214. doi: 10.1186/s13059-021-02419-7
51. Adams AK, Kristy BD, Gorman M, Balint-Kurti P, Yencho GC, Olukolu BA. Qmatey: an automated pipeline for fast exact matching-based alignment and strain-level taxonomic binning and profiling of metagenomes. Brief Bioinform. 2023;24(6). doi: 10.1093/bib/bbad351
52. Blin K. ncbi-genome-download Zenodo2023 [Available from: https://zenodo.org/records/8192486.
53. Bushnell B, editor BBMap: A Fast, Accurate, Splice-Aware Aligner2014.
